# Complementary contributions of dorsal and ventral striatum to cost-benefit vigor adaptations

**DOI:** 10.1101/2024.05.31.596850

**Authors:** Thomas Morvan, Zelda Timmel, Christophe Eloy, David Robbe

## Abstract

Humans and other animals adjust when and how fast they execute goal-directed actions according to expected costs and benefits. Yet, the behavioral principles and brain regions underlying adaptive vigor control remain poorly understood. Here, we developed a self-paced foraging task in which rats must run across a motorized treadmill to collect rewards, allowing separate manipulation of benefit (reward delivery probability) and cost (effort). Throughout the foraging sessions, run timing and speed jointly reflected the rats’ idiosyncratic urgency to obtain rewards. In contrast, when the cost of crossing the treadmill increased, rats ran faster to maintain a high reward rate while run timing remained stable. Consistent with distinct mechanisms for adapting to costs or benefits, dorsal striatum lesions primarily limited running speed under effortful conditions, whereas ventral striatum lesions reduced reward-seeking urgency at session onset. Together, these findings suggest that vigor is modulated by costs and benefits through distinct striatal circuits.

## Introduction

In a wide range of contexts, humans and other animals adjust simultaneously the timing and speed of their actions according to expected benefits and costs. For instance, when a mouse detects a looming threat, it quickly initiates a fast run toward the closest shelter. Similarly, to feed their hungry chicks, birds will make numerous and rapid back-and-forth flights between their nest and a rich food source. These modulations of vigor, defined here as the timing and speed of actions, reflect the tendency of animals to maximize their capture rate (CR), i.e., the sum of acquired rewards (or reached goals) minus the total effort expended (or cost of moving), divided by the total time invested (CR = (Rewards − Efforts)/Time, [1–3]). Animals therefore behave according to the utility of the options available to them, utility that depends not only on expected reward and effort but also on the passage of time. Indeed, time discounts the value of rewards [4] and conceptualizing it as a cost explains why it is advantageous to quickly initiate and execute goal-oriented actions [1, 5–9]. Still, hasty decisions can be maladaptive, and vigorous actions are prone to inaccuracy [2], highlighting the need for mechanisms that jointly regulate the timing and the speed of goal-oriented behaviors.

To provide quantitative evidence for the joint influence of utility on the timing and speed of voluntary actions, the co-variation between reaction time and movement duration has been studied in motor tasks with varying reward or effort levels. This revealed that both reaction time and movement duration were shorter when humans reached for targets associated with rewards [10]. Conversely, reaction time increases when subjects have to perform longer (i.e., more costly) movements [11]. Moreover, in a gaze foraging task, human subjects took longer to initiate a new saccade toward another image after a series of trials with low reward outcomes, and executed this saccade more slowly [12]. Finally, instructing subjects to move quickly is enough to reduce their reaction time [13]. To account for these co-modulations of action timing (when to act) and speed (how fast to act) by utility, a shared neuronal mechanism has been proposed, potentially involving the basal ganglia [2, 3, 14].

A role of the basal ganglia in the coordination of action timing and speed is supported by the well-known behavioral deficits associated with their dysfunctions [15] and their role in cost-benefit decision-making and motor control [16, 17]. Both dorsal (i.e., associative and sensorimotor) and ventral (i.e., limbic) regions of the basal ganglia have been implicated in vigor control in humans, with potentially complementary functions. Dorsal regions have been suggested to contribute to scaling movement speed according to effort requirement [18–21], whereas ventral ones track the value of rewards [22, 23] and invigorate behavior accordingly [24]. In rodents, the dorsal and ventral striatum (DS, VS) have also been implicated in vigor control, both through perturbation approaches and by identifying neurophysiological correlates of movement kinematics [25–41]. Surprisingly, these regions are often assumed to subserve orthogonal functions, with DS being linked to action selection, motor control and procedural learning [42–46], and VS to reward processing and motivation [47–51]. Although a primary role of the sensorimotor regions of the basal ganglia in the control of movement vigor has been proposed [52, 53], its underlying computational principles remain elusive in rodents. In particular, the factors thought to shape movement vigor, such as expected reward, effort, and urgency, are more difficult to dissociate than in humans and non-human primates [14, 54], leaving the respective contributions of DS and VS largely undefined.

Here, we developed a self-paced foraging task in which rats freely modulate the timing and speed of short running bouts to collect rewards delivered at the extremities of a motorized treadmill. Benefits (probability of obtaining reward) were manipulated within sessions, whereas motor cost (length or velocity of treadmill) varied between sessions. We used a model-based approach to capture how rats adapted the timing and speed of their reward-oriented runs during and across sessions. This framework revealed that, as sessions progressed, the animals’ urgency to obtain rewards was reflected in both run timing and speed. In contrast, when the motor cost associated with crossing the treadmill increased, all rats ran faster to preserve their reward rate, without marked alteration of run initiation timing. Consistent with distinct mechanisms for adapting vigor to benefits or costs, ventral striatum lesions reduced the urgency to obtain water at session onset, while dorsal striatum lesions primarily limited the rats’ tendency to run faster under high-cost conditions.

## Results

### Model-based quantification of run timing and speed adaptations during self-paced foraging in freely moving rats

To better understand how costs and benefits influence the timing and speed of reward-oriented actions in freely moving animals, we developed a self-paced foraging task in which water-restricted rats (Methods) could run between two platforms located at the extremities of a motorized treadmill, each equipped with a spout delivering calibrated drops of water (Methods). Upon reaching a platform, a single reward was delivered according to a probabilistic rule and rats had to cross back to the opposite platform to receive the next one (Fig. 1A, left). Rats learned the principle of this task in a matter of a few sessions (Methods). Their behavior was characterised by straight runs between the two platforms interleaved with pauses of varying durations, including a very brief licking bout, allowing us to separate runs from inter-run periods (Methods, Fig. 1A, right). In the rest of the manuscript, inter-run durations will serve as a proxy for run initiation times, also referred to as run timing, and run durations as a proxy of running speeds.

**Figure 1:**
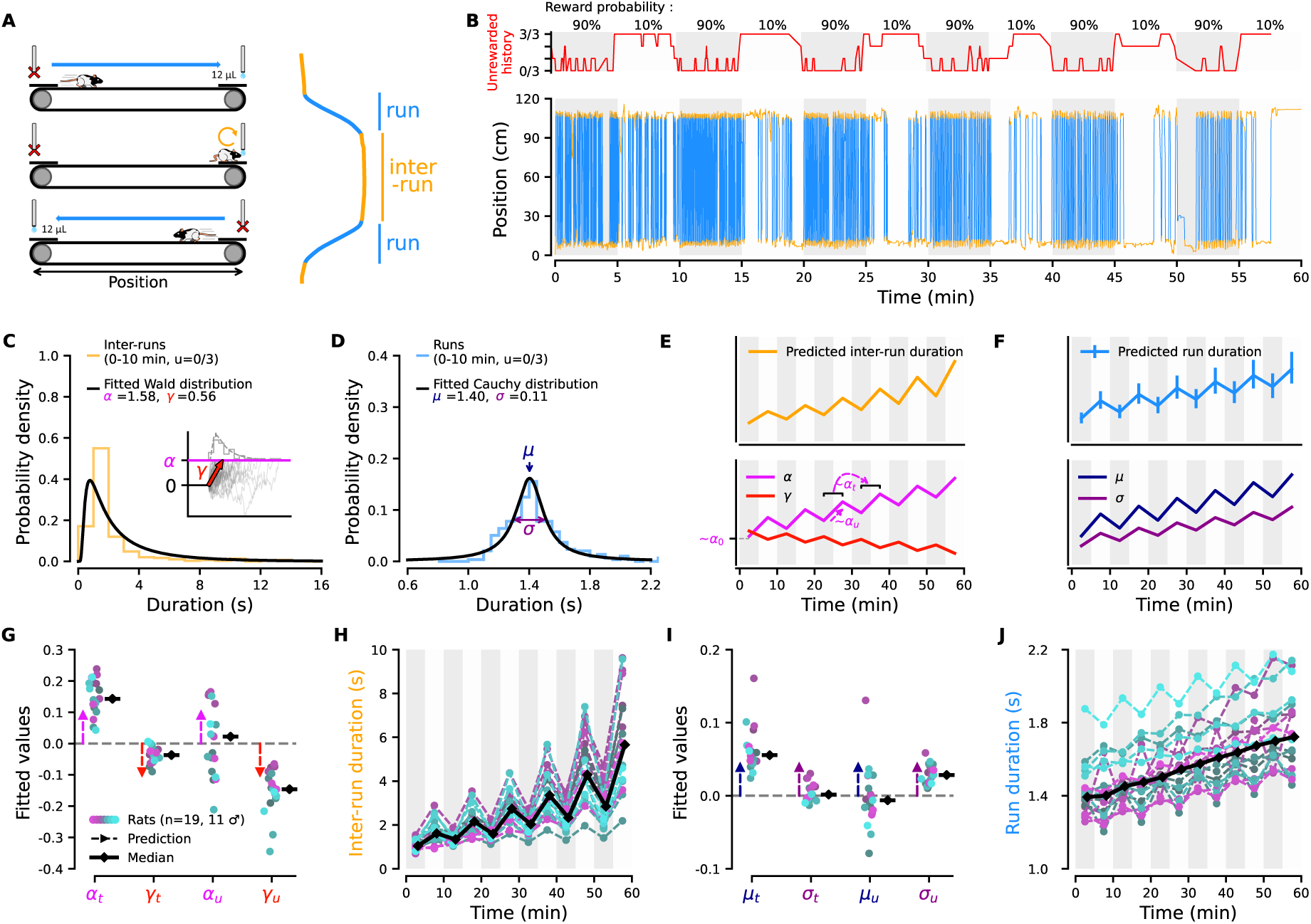
Run timing and speed are co-modulated by individually distinct perceptions of foraging utility. **A**, Left, task schematic. Right, trajectories are separated into run and inter-run epochs. **B**, Example session (longest distance, stationary treadmill). Top, reward probability alternated between high (90%, gray) and low (10%, white) in uncued 5-min blocks. The red line shows the ratio of unrewarded runs over the last 3 runs. Bottom, rat trajectory over time (same color code as A) overlaid on reward probability blocks. **C**, Inter-run durations distribution from the first 10 min of sessions (same animal and condition as B, unrewarded history *u* = 0/3, orange) and fitted Wald distribution (black) with parameters α and γ. Inset, drift-diffusion model with α as threshold (magenta) and γ as drift rate (red). Gray histogram and dashed line show simulated and fitted distributions with the same parameters. **D**, Run durations distribution (blue, same condition as C) and fitted Cauchy distribution (black) with median µ and scale σ. **E**, Top, utility-based prediction of inter-run duration modulations across reward probability blocks and time. Bottom, corresponding modulation of α and γ. α_0_, α_t_, and α_u_ capture, respectively, baseline α, its modulation by time, and by unrewarded history. **F**, Top, similar to E for run durations, µ and σ (bottom). **G**, Expected (arrows) and observed (dots) modulation of α and γ by time and unrewarded history. **H**, Inter-run duration modulation across rats (same motor cost as B). Colored lines: per-animal median reconstructed from α(*t*, *u*) and γ(*t*, *u*) (Methods); black line: group median. **I**, Similar to G for µ and σ. **J**, Similar to H for run duration modulation reconstructed from µ(*t*, *u*) and σ(*t*, *u*).

To examine how the timing and speed of the runs were modulated by fluctuations in expected benefits, the probability of obtaining a drop of water upon reaching a platform was manipulated in unsignaled blocks of 5 minutes during one hour-long sessions (rats performed two sessions per day), alternating between high (90 %) and low (10 %) (Fig. 1B, top, alternating gray and white blocks respectively). To examine the influence of motor cost on run timing and speed, we used two types of manipulations. Every day, we changed either the distance between platforms (29 cm, 62 cm, or 94 cm) while the treadmill belt was immobile, or the belt velocity (20, 10, ±2, −10, or −20 cm/s relative to the running direction) in the long-distance configuration. Rats performed 6 sessions in each of these eight motor cost conditions (i.e., 3 days per condition). They were first tested in the 3 distance conditions (9 days of distance manipulations, pseudorandom order conserved across animals). Then, after being habituated to the moving belt, they were tested with the 5 belt velocities (15 days of treadmill velocity manipulations, pseudorandom order conserved across animals; see Methods).

We first examined how run timing and speed were modulated during foraging sessions in which the distance between platforms was the longest (i.e., 94 cm) and the treadmill belt was immobile. As expected, a representative session from a single animal showed fewer runs during low-reward probability blocks (Fig. 1B, bottom, light gray blocks). In addition, the number of runs declined over the course of the session, consistent with a progressive reduction in motivation, possibly due to satiety, fatigue, or both. Such a reduction in the number of runs renders direct within-session comparisons of their timing and speed difficult. To mitigate this issue, for each rat, run and inter-run durations from sessions performed under the same motor cost condition were pooled together and divided into 6 bins of 10 minutes. Each bin was further divided according to the number of unrewarded runs in the preceding three runs (unrewarded history score: 0/3, 1/3, 2/3, 3/3; Fig. 1B, top red line; see Methods). Visualizing one of the 24 resulting inter-run duration distributions shows that it is dominated by very small values and displays a long tail reflecting rare, long events (Fig. 1C, orange histogram, same animal as in Fig. 1B). Thus, rats produce runs organized in bursts with short intra-burst pauses and a few much longer inter-burst pauses, yielding a distribution that cannot be adequately summarized by its mean and variance. This motivated the use of the Wald distribution, whose two parameters, α and γ, can provide an accurate characterization of the different patterns of inter-run durations (Fig. 1C, black line; Methods). For example, an increase in the prevalence of long pauses may occur with little change in the median, yet substantially alter the shape of the distribution. Such changes are naturally captured by variations in α and γ. Incidentally, these parameters correspond directly to the threshold height and drift rate parameters, respectively, of drift-diffusion models (DDMs), which have been widely used to account for reaction-time and decision-time distributions across a broad range of behavioral tasks [55–58] (Fig. 1C, inset). Importantly, by quantifying inter-run durations with DDM-related summary statistics, we do not imply that these events correspond to periods of explicit deliberation about when to run again, during which animals remain immobile. Rather, the DDM correspondence simply helps interpret how changes in α and γ shape the distribution of inter-run durations, and, by extension, how the timing of the runs is modulated throughout the foraging sessions. For instance, an increase in the threshold α combined with a decrease in the drift rate γ would account for both a rightward shift of the inter-run duration distribution peak and an increase in rare, long pauses. Conversely, a decrease in both α and γ would lead to a leftward shift (or no shift) of the distribution peak, together with a higher number of long pauses (i.e., maintained or increased intra-burst frequency, but decreased inter-burst frequency). Finally, to capture the intra-session modulations of running speed, the same partitioning was performed on run durations (i.e., according to time and unrewarded history). The 24 run duration distributions obtained (e.g. Fig. 1D, blue histogram) were well captured by a Cauchy distribution with median µ and spread parameter σ, the half-width at half-maximum of the distribution (Fig. 1D, black line, Methods).

The timing and speed of reward-oriented actions have been shown to reflect their utility [3]. Accordingly, simple predictions can be made regarding the modulation of inter-run and run durations during foraging sessions. Inter-run durations are expected to increase both when animals transition from high- to low-reward probability blocks and over the course of the session as animals become satiated and fatigued (Fig. 1E, top). In terms of the Wald distribution parameters, these predictions translate into a sawtooth-like increase in α (the DDM threshold) and/or decrease in γ (the drift rate) over the course of the session (Fig. 1E, bottom). If, in our experimental condition, run timing and speed are co-modulated by utility, run durations should exhibit a similar pattern of modulation over the course of the session (Fig. 1F, top), resulting in an increase in µ during low-reward probability blocks and as the session progresses (Fig. 1F, bottom, dark blue line). Moreover, as low-reward conditions increase movement variability in both humans and rodents [59, 60], running speed variability σ is expected to increase during low-reward probability blocks and over the course of the session (Fig. 1F, bottom, dark magenta line).

To capture within-session modulations of run initiation times, we first fitted a Wald distribution to each of the 24 inter-run duration distributions (obtained by dividing sessions according to time and unrewarded history), leaving α and γ as free parameters. We then fitted the resulting 24 estimates of α and γ with the following linear model, capturing their baseline values and modulation by session time and unrewarded history:

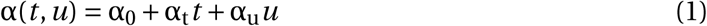

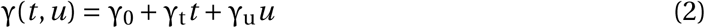

with α_0_ and γ_0_ describing the inter-run durations distribution at the beginning of the sessions, α_t_ and γ_t_ describing its linear modulation over time *t*, and α_u_ and γ_u_ describing its modulation by the history of unrewarded trials *u* (see annotation on Fig. 1E, bottom, for an approximate representation of α_0_, α_t_, and α_u_).

The same two-step approach was applied to run durations, fitting a Cauchy distribution to extract µ and σ, then modeling their session-wise modulations as:

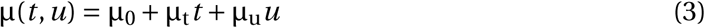

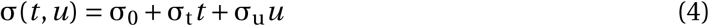

with µ_0_ and σ_0_ describing the initial state of the run durations distribution at the beginning of the sessions, µ_t_ and σ_t_ describing its modulation over time *t*, and µ_u_ and σ_u_ describing its modulation by the unrewarded history *u*.

This type of linear model capitalizes on all available data, reducing estimation errors in sparse blocks by leveraging information from the rest of the session. Goodness-of-fit analyses for both inter-run and run durations indicated that the model provided an accurate description of the data (Supplementary S1; Methods). Furthermore, model-based parameters correlated strongly with model-free estimates (Supplementary S2; Methods), and the inferred session-wise dynamics closely reproduced those obtained using model-free quantification (compare Fig. 1H,J with Supplementary S2C,F). Finally, comparisons of α*_u_*, γ*_u_*, µ*_u_*, and σ*_u_* between the first and second halves of the session revealed no significant differences, indicating that the effect of reward history on run timing and speed remained stable throughout the session (Supplementary S3).

Using this model-based quantification, we examined whether session-wise change in run timing and speed in 19 rats (11 males, blue side of the gradient, color code of individual animals maintained across plots) followed utility-based predictions (Fig. 1G,I; dashed arrows indicate the predicted direction based on Fig. 1E,F, bottom). As predicted, α increased and γ decreased over the course of the session (Fig. 1G; α_t_ > 0, one-sided Wilcoxon signed-rank test, *p* = 1.9×10^−6^; γ_t_ < 0, one-sided Wilcoxon signed-rank test, *p* = 1.3×10^−5^), consistent with the observed global reduction in run frequency and the emergence of longer pauses as the session progressed (Fig. 1B, bottom). In contrast to the systematic effects of time, the influence of unrewarded history on run timing was unexpectedly heterogeneous across animals, the coefficient α_u_ being positive in some animals and negative in others (Fig. 1G; two-sided Wilcoxon signed-rank test, *p* = 0.74). On the other hand, γ was systematically decreased by unrewarded outcomes (Fig. 1G; γ_u_ < 0, one-sided Wilcoxon signed-rank test, *p* = 1.9 × 10^−6^), and animals with the most negative α_u_ showed a greater decrease in γ (α_u_ correlated with γ_u_, Supplementary S4A). This correlation accounts for the systematic presence of long pauses during low-reward probability blocks across all animals (Fig. 1H). Critically, rats with the most negative α_u_ showed limited change in their average inter-run duration between high and low reward probability blocks (Supplementary S4B, C), indicating that their intra-burst run frequency, the main determinant of their average run frequency, was preserved. Overall, the sawtooth-like increase in average inter-run duration through the session is congruent with utility-based predictions on action timing (compare Fig. 1H with Fig. 1E, top; Methods). However, our model-based approach revealed a subset of rats that maintained short inter-run durations together with rare long pauses during low reward probability blocks, potentially reflecting increased urgency to obtain water when they faced an unexpected decrease in reward outcomes [61, 62].

Next, we examined across animals how the parameters describing run duration distributions were modulated during the sessions. As predicted, µ consistently increased over time (Fig. 1I, µ_t_ > 0, one-sided Wilcoxon signed-rank test, *p* = 1.9 × 10^−6^) indicating that run duration, like inter-run duration, progressively increased over the course of the sessions. Run duration variability, on the other hand, remained stable (σ_t_ = 0, two-sided Wilcoxon signed-rank test, *p* = 0.47). In contrast to utility-based predictions, µ_u_ displayed striking interindividual differences, being positive in some animals (decreased running speed) and negative in others (increased running speed; Fig. 1I; two-sided Wilcoxon signed-rank test against zero, *p* = 0.59), echoing the pattern observed for α_u_. On the other hand, σ systematically increased with unrewarded history (Fig. 1I, σ_u_ > 0, one-sided Wilcoxon signed-rank test against zero, *p* = 1.9 × 10^−6^).

Comparing the average evolution of the inter-run and run durations (Fig. 1H and J) revealed a more subtle picture than thought from the expected co-modulation of action timing and speed by utility. Indeed, if both inter-run and run durations increased with session time, their modulation by the high and low reward opportunity blocks appeared distinct (compare Fig. 1H with Fig. 1J; note the absence of sawtooth pattern in the group median of the latter). Still, several animals displayed a modulation, positive or negative, of their run duration by the high and low reward-probability blocks (see sawtooth pattern of individual dashed lines in Fig. 1J, and the positive and negative individual values of µ_u_ in Fig. 1I). Interestingly, α_u_ (the modulation of the run initiation threshold by unrewarded outcomes) also displayed positive and negative individual values. In fact, a comprehensive pairwise correlation of behavioral parameters confirmed that α correlated strongly with µ, with α_t_ and α_u_ correlating with µ_t_ and µ_u_, respectively (Supplementary S4D,E; Methods). In contrast, no significant correlations were observed between the drift rate γ and µ (Supplementary S4F). Because animals with the most negative α_u_ (i.e., with the strongest reduction µ) displayed preserved high intra-burst frequency (Supplementary S4B, C), changes in perceived utility (driven by satiety or idiosyncratic reaction to unrewarded outcomes) appear to modulate running speed and intra-burst frequency in a coupled manner, while affecting the inter-burst longer pauses independently. This result validates the use of the Wald distribution, showing that its two parameters capture non-redundant aspects of behavior.

### Rats adapt to increasing motor cost by running faster

A key feature of our task is that the motor cost required to cross the treadmill can be manipulated separately from reward opportunity. We examined how the timing and speed of runs varied across motor cost conditions, focusing on the baseline values derived from the linear models describing their session-wise modulation (i.e., α_0_, γ_0_, µ_0_ and σ_0_). According to utility-based predictions, increasing motor cost should delay run initiation, which would translate into a higher threshold α_0_, and/or a lower accumulation rate γ_0_ (Fig. 2A, left). It should also lead to longer and more variable runs (higher µ_0_ and σ_0_, Fig. 2A, right). We modulated the motor cost by changing, across sessions, the distance (d) between the platforms (Fig. 2B, left) or the treadmill belt’s velocity (*v*_belt_, Fig. 2B, right). Contrary to the above predictions, α_0_ and γ_0_ remained unchanged across motor cost conditions (Fig. 2C, D). On the other hand, increasing the distance between the platforms led to a significant increase in the baseline run duration µ_0_ (Fig. 2E, left). Importantly, this increase was much smaller than expected from a one-to-one relationship between motor cost and run duration (compare median values with the expected unity line, dashed orange). Indeed, rats run faster when the distance between platforms increased from 29 cm to 94 cm (Fig. 2F, left). Manipulations of treadmill velocity revealed a similar pattern: run durations were largely stable across velocities, increasing only when the treadmill maximally opposed the animals’ movement (Fig. 2E, right). This stability implies that rats ran faster when the treadmill opposed their movement and slower when it facilitated it, as confirmed by each animal’s running speed across motor cost conditions (Fig. 2F, right, Supplementary Video). Finally, σ_0_ increased when motor cost became very high, consistent with a utility-based modulation of speed variability (Fig. 2G). Together, these findings show that varying motor costs robustly modulate kinematic parameters of reward-oriented runs, while largely sparing run timing.

**Figure 2:**
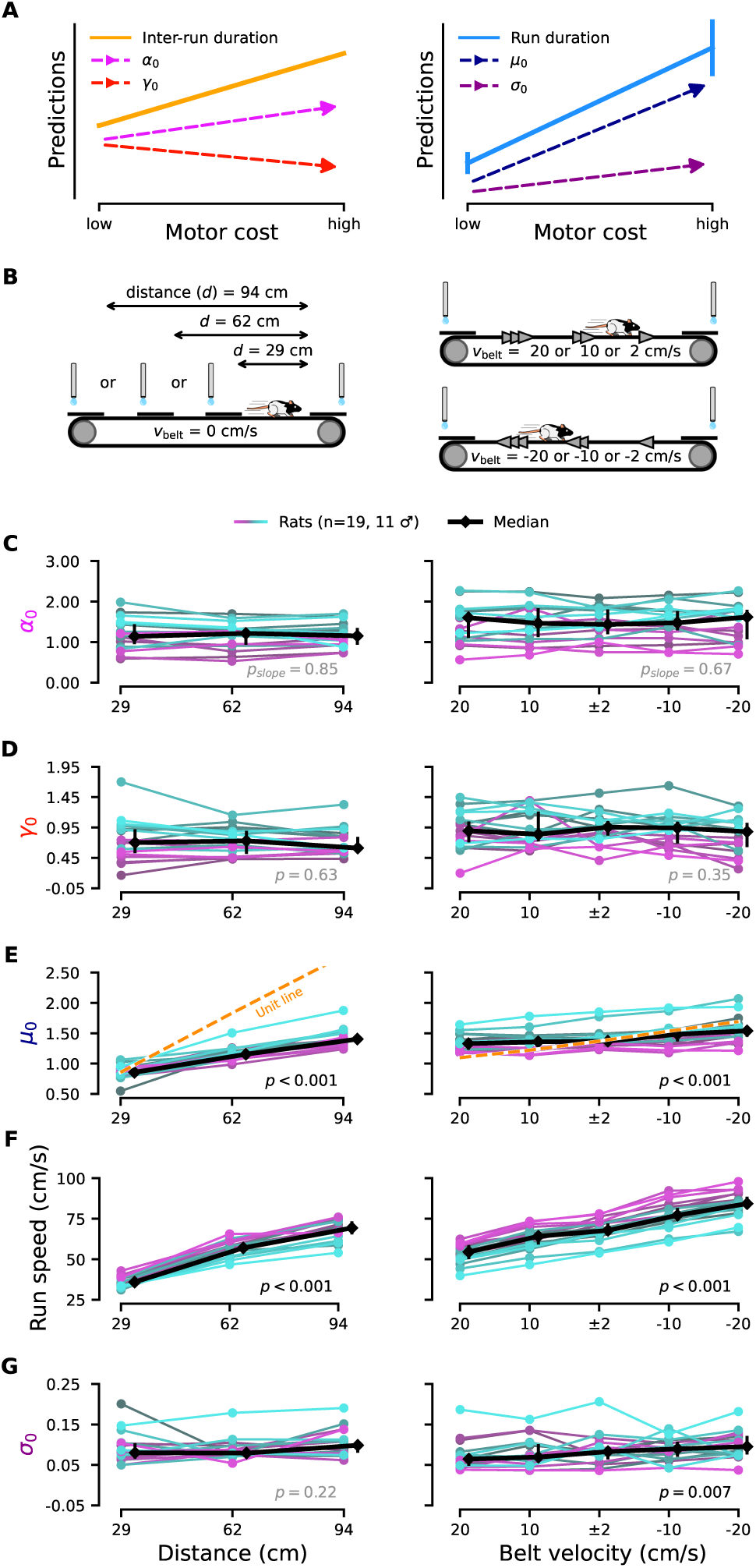
Motor cost selectively modulates running speed while sparing run timing. **A**, Utility-based predictions for run timing (left) and run duration (right) with increasing motor cost, and associated changes in model parameters (same color code as Fig. 1E,F). **B**, Motor cost manipulations by changing platform distance (left) or belt velocity (*v*_belt_, right, Methods). **C-G**, Effect of distance (left) and belt velocity (right) on α_0_ **(C)**, γ_0_ **(D)**, µ_0_ **(E)**, running speed **(F)**, and σ_0_ **(G)**. In E, the unity line indicates the expected run duration at constant speed from the 29 cm (left) or ±2 cm/s (right) condition. Black, population median; error bars, interquartile range. Significance of slopes (*p*-values) assessed with permutation test (Methods).

So far, our results show a utility-driven covariation of run timing and speed during the sessions (significant correlations between α_u_ and µ_u_, and between α_t_ and µ_t_) which is not hardwired, as rats responded to increasing motor costs by selectively modulating their running behavior. This suggests that, in our experiments, µ is sensitive to both the motor cost and the benefit associated with running between platforms, whereas α is primarily sensitive to benefit. To directly test this possibility, we sought to disentangle the respective contributions of reward and effort to α_t_ and µ_t_. Typically, effort and satiety are perfectly correlated, precluding such an analysis. However, in our experimental setting, animals tended to maintain short run durations across the eight motor cost conditions and continued running in low-probability blocks, resulting in a situation in which the total number of rewards collected was not perfectly correlated with effort expended (Supplementary S5A,B; E_TOT_ and R_TOT_, ρ = −0.36, see Methods).

We therefore used linear mixed models to quantify whether session-to-session variation in α_t_ and µ_t_ could be predicted by total effort per session (E_TOT_) or total reward obtained (R_TOT_, see Methods for details). This analysis confirmed that µ_t_ was sensitive to both benefit and cost, whereas α_t_ was primarily driven by reward (Supplementary S5C-F), reinforcing the idea that, during foraging, costs and benefits are integrated via distinct mechanisms.

### Run duration minimizes the costs of movement and time

The observation that rats ran slower (faster) when the treadmill facilitated (impeded) their run toward the platforms suggests that they balance effort and cost of time to maintain a nearly constant crossing duration (Fig. 2F, right). The presence of interindividual differences prompted us to develop a simple model to capture how each animal modulated its running speed across motor cost conditions. A parsimonious interpretation of the aforementioned modulations is that baseline run duration, µ_0_, reflects the minimization of both movement and time costs [63, 9, 64]. Because the energetic cost of movement increases with locomotion speed [65–67], we examined the animals’ speed profiles as they crossed the treadmill and found that their shape was conserved across motor cost conditions (Fig. 3A). Thus, they can be normalised (Fig. 3B, individual lines) and formally described using a scaled universal function that depends on V_max_, the maximum speed reached by the animal and T, the run duration (Fig. 3B, inset, see Methods). This universal function allowed us to estimate the cost of movement C_M_ in a relatively simple manner as follows:

**Figure 3:**
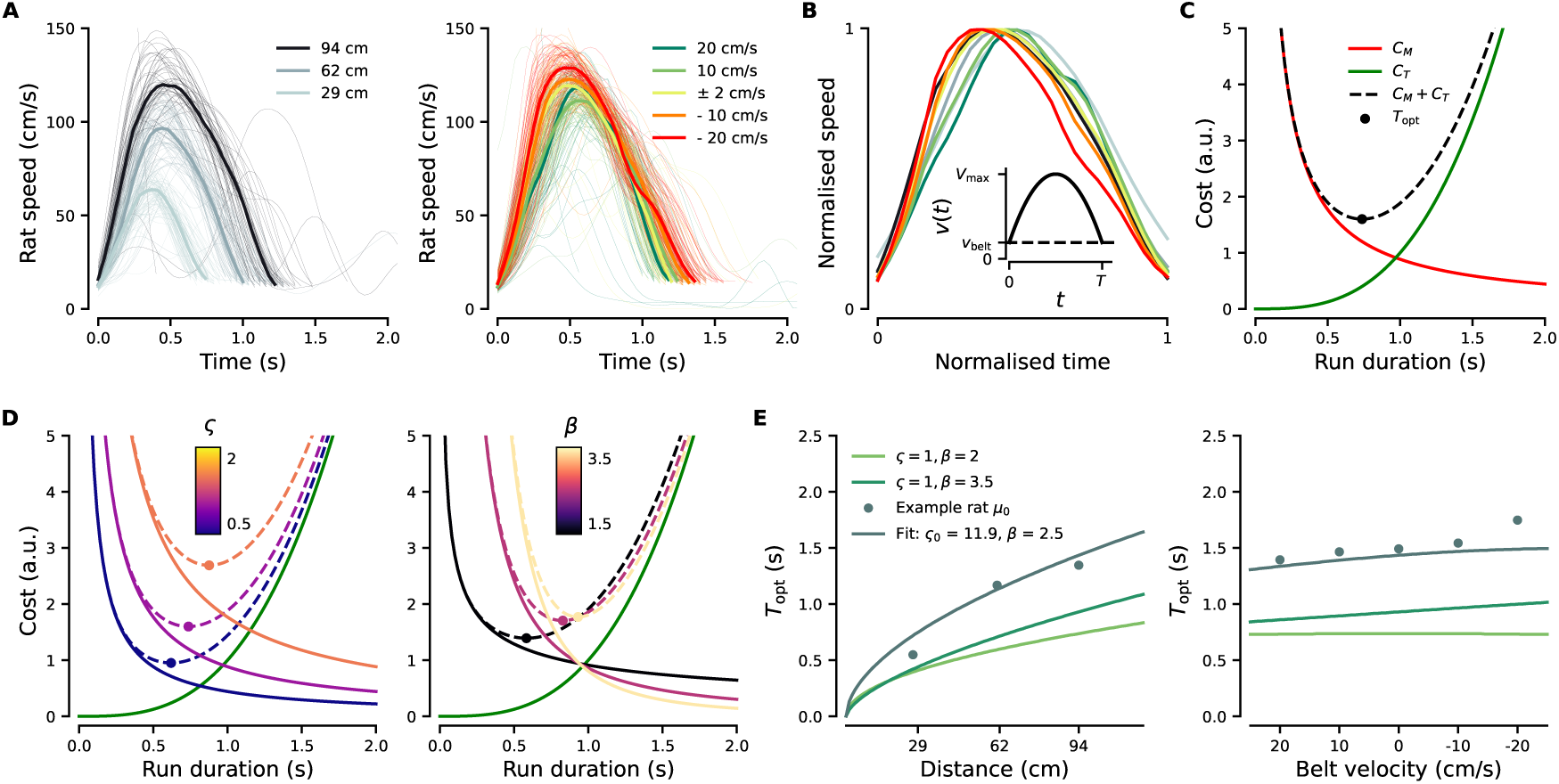
Run duration minimizes the costs of movement and time. **A**, Running speed profiles across distances (left) and treadmill belt velocities (right). **B**, Rescaling speed and time by V_max_ and T reveals a universal speed profile (Inset). **C**, The optimal run duration T_opt_ minimizes the total cost C that sums the costs of movement (C_M_) and time (C_T_). **D**, Effects of ς (the relative sensitivity to the costs of movement and time, Equation M14, Middle) and β (the exponent linking running speed to movement cost, Equation M11) on T_opt_. **E**, Predicted optimal run durations across motor costs using the cost minimization model with fixed ς and β (green lines, Methods). Teal dots show µ_0_ of the example animal; the teal line shows the model prediction with fitted ς_0_ and β values.

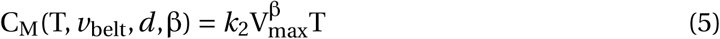

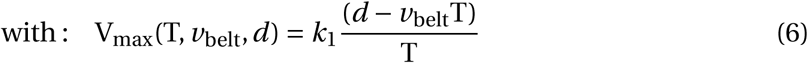

C_M_ depends on V_max_ and T. V_max_ can be derived from the distance traveled *d*, the run duration T and the velocity of the treadmill belt *v*_belt_; *k*_1_ and *k*_2_ are two constants that only depend on the shape of the universal function. Critically, C_M_ increases with V_max_ to the power β, with β a tunable parameter. In our model, we considered that the optimal duration of a run minimizes a cost function C which is the sum of the cost of movement C_M_ and a cost linked to the passage of time C_T_ (Fig. 3C, red, and green lines, respectively, see Methods). Because animals could be more or less sensitive to one of these two costs, the cost function C can be expressed as:

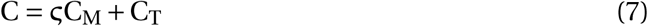

with ς a tunable parameter indicating the relative sensitivity to the movement cost C_M_ against the cost of time C_T_. As expected, increasing ς or β results in longer optimal run durations (Fig. 3D; note that all parameters are fixed to 1 in this illustration to show their independent effects on model behavior; fitted values for one animal are shown in Fig. 3E). We then predicted the optimal run durations as a function of platform distance and treadmill belt velocity using the cost function C, for different values of β with a fixed ς (Fig. 3E, light and dark green lines). The observed increase in run duration with distance is similar to our experimental observations (Fig. 3E, left). Conversely (and trivially), this cost minimization model also reproduces the fact that the animals’ running speed increased with distance and displayed a linear relationship with the belt velocity (Fig. 3E, right). Finally, we found for each animal the values of ς_0_, ς_t_, ς_u_ and β that provided the best fit to the run duration model parameters µ_0_, µ_t_, µ_u_ across motor costs (Fig. 3E, teal dots correspond to the µ_0_ obtained for the example animal, teal line corresponds to the run duration predicted by the cost minimization model, see Methods). Altogether, our cost-minimization model provides simple metrics (ς and β) to quantify how each animal adjusts its running speed across motor-cost conditions and over the course of a session. In addition, the model offers a mechanistic intuition for these behavioral modulations by linking changes in run duration to the minimization of movement costs (Supplementary S6; see Methods).

### Distinct alterations of vigor following lesions of dorsal or ventral striatum

So far, our experiments and model-based analyses reveal that reward-related alterations in the utility to run affect both the timing and speed of reward-oriented movements (Fig. 1). In contrast, when motor cost was manipulated, animals ran faster to save time, whereas run timing remained largely unaffected (Fig. 2). Together, these findings raise the possibility that distinct mechanisms translate perceived costs and benefits into adaptive behavior. The dorsal and ventral regions of the striatum are well positioned to support this functional dissociation. Indeed, indirect evidence suggests that the dorsal striatum (DS) regulates the speed of reward-oriented movements based on estimated motor costs [18, 19, 35, 20, 68, 69], whereas the ventral striatum (VS) plays a central role in reward processing [70, 24]. To investigate this possibility, we performed excitotoxic lesions of neurons in either the dorsal striatum (DS; *n* = 19, 9 females) or the ventral striatum (VS; *n* = 15, 7 females; Fig. 4A). To avoid confounds related to the repetition of our long experimental protocol, particularly the pronounced weight gain in male rats, lesions were performed prior to behavioral experiments; non-lesioned animals served as controls rather than a sham-lesion group (Methods). We first compared the average modulation of run timing and speed in the long-distance / treadmill-off motor cost condition (as in Fig. 1), in DS-, VS-lesioned and non-lesioned (control) animals. In both lesion groups, run timing remained sensitive to the alternation between high- and low-reward probability blocks (Fig. 4B). However, while controls and DS-lesioned animals exhibited short inter-run durations at the beginning of the session that progressively increased (Fig. 4B, top), this early-session urgency was markedly attenuated in VS-lesioned animals. They began each session with longer inter-run durations that remained elevated throughout (Fig. 4B, bottom). Regarding running speed, both DS- and VS-lesioned rats displayed longer run durations than controls (Fig. 4C). Notably, DS-lesioned animals, like controls, showed a progressive increase in run duration over the course of the session, albeit shifted toward overall longer durations (Fig. 4C, top). VS-lesioned rats did not show the session-wise progressive increase in run duration; instead, as observed for inter-run durations, they began the session with longer run durations that remained elevated throughout (Fig. 4C, bottom).

**Figure 4:**
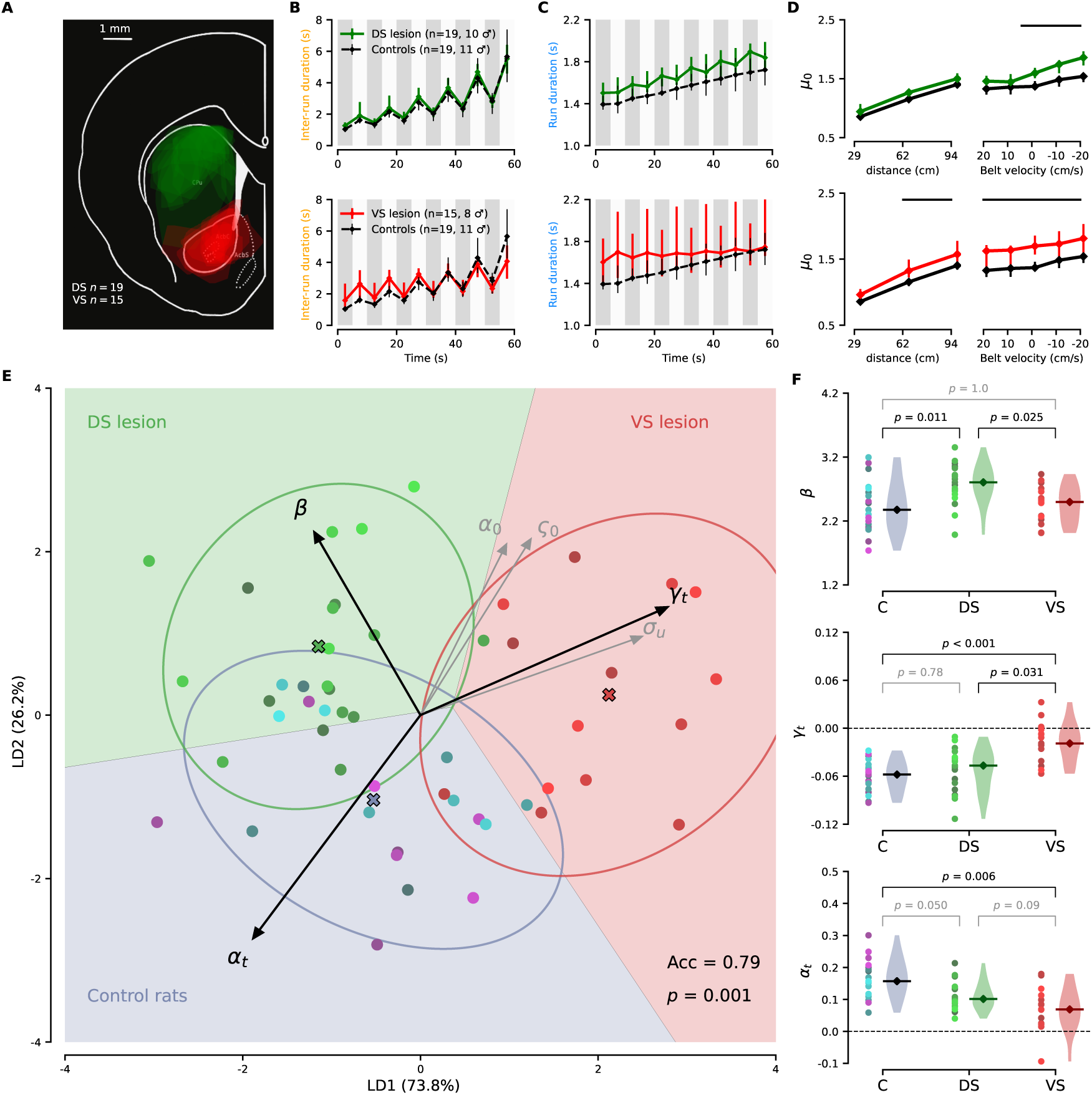
Dorsal and ventral striatum as distinct substrates for cost-driven speed and motivational urgency. **A**, Lesion outlines (dStr, green, *n* = 19; vStr, red, *n* = 15; Methods) superimposed on a coronal slice adapted from the Paxinos rat brain atlas. **B**, Inter-run durations for control (dashed black line) and lesioned rats (top, dStr, green; bottom, vStr, red). Points represent block medians reconstructed from α(*t*, *u*) and γ(*t*, *u*) (same as Fig. 1H); error bars indicate interquartile range. **C**, Similar to B for run durations. **D**, Median µ_0_ across the 8 motor cost conditions for control (black) and lesioned rats (top, dStr, green; bottom, vStr, red). Error bars show interquartile range. Significant differences between group medians assessed by permutation test (black horizontal lines, Methods). **E**, Model parameters projected onto the first two linear discriminants (LD1 and LD2; percentage of explained variance on each axis). Each point represents one animal (control, blue; dStr, green; vStr, red); crosses indicate group centroids; ellipses denote ±2 standard-deviation group dispersion. Shaded regions show classifier decision boundaries. Arrows show variable loadings (length proportional to contribution); only variables with significant inter-group differences are shown; the three strongest contributors are in black. Acc = accuracy, *p*-values from permutation test with group labels (Methods) **F**, Inter-group differences for the three parameters with the strongest loadings: β, γ_t_, and α_t_. *p*-values from permutation test defined as the fraction of shuffles with a median difference greater than observed; Bonferroni correction applied.

Next, we examined how DS- and VS-lesioned animals modulated their running speed across motor cost conditions. Consistent with an overall reduction in vigor, VS-lesioned animals ran more slowly than control rats across most motor cost conditions, as reflected by their longer running durations, including in conditions in which the treadmill belt facilitated movement toward the platform (Fig. 4D, bottom). In contrast, DS-lesioned animals displayed progressively longer run durations as motor cost increased (Fig. 4D, top), indicating a reduced tendency to increase running speed when task demands required it. Importantly, this effect occurred while the speed profile was mainly preserved, with DS-lesioned rats only displaying a lower peak speed under the highest motor cost (Supplementary Fig. S7A-D), ruling out major locomotor impairment as a source of increased run duration. In the lowest motor cost condition, speed distributions were indistinguishable between groups (Supplementary Fig. S7A,E,F). In the highest motor cost condition, the distribution was shifted toward lower speeds in DS-lesioned animals; however, the highest speeds reached did not differ significantly between groups (Supplementary Fig. S7G,H). Together, these results indicate that DS-lesioned animals retain the capacity to run fast but express high running speeds less frequently when task demands require it.

Next, we leveraged our model-based approach to further investigate whether DS- and VS-lesioned rats exhibited distinct profiles of vigor alterations. To provide an integrated view of pairwise comparisons across the three groups for all model-derived parameters, we used linear discriminant analysis (LDA), an approach that captures multivariate group differences in a low-dimensional space (see Methods). Projection of each animal onto the first two linear discriminant axes revealed a clear and significant separation among the three groups, driven by a limited subset of parameters (Fig. 4E). DS-lesioned animals primarily differed from control and VS-lesioned animals by their increased β values, which are inversely proportional to the propensity to increase running speed in response to higher motor costs (Fig. 4E, 4F, top). On the other hand, animals with VS lesions differed from the other groups by a reduced modulation of γ by session time (Fig. 4E, 4F, middle). Together with the markedly attenuated α_t_ (Fig. 4F, bottom), this pattern accounts for the overall stability of run timing in this group, consistent with a diminished urgency to produce many runs at the beginning of the session. Both lesioned groups showed an increased baseline relative sensitivity to time versus effort (ς_0_; Fig. 4E; Supplementary S8D), consistent with DS-lesioned animals being more sensitive to effort and VS-lesioned animals less sensitive to time. As ς_0_ and β increased with body weight in non-lesioned animals (Supplementary S8F,G), we verified that the three groups did not differ in their weight (Control: median = 253.2 g, IQR = [208.1–271.8], *n* = 19; DS-lesioned: median = 224.8 g, IQR = [199.3–270.6], *n* = 19; VS-lesioned: median = 250.0 g, IQR = [197.5–282.5], *n* = 15; all pairwise permutation tests: *p* > 0.05 after Bonferroni correction). Altogether, these results show that while both DS- and VS-lesioned animals exhibited reduced vigor and shared alterations of a few parameters (Supplementary S8 and see Discussion), their deficits were also qualitatively distinct: DS lesions impaired the adaptation of running speed to motor costs, whereas VS lesions primarily disrupted the motivational drive to make frequent and fast runs at the beginning of the session.

## Discussion

In this study, we used a self-paced foraging task for freely-moving rats to understand how changes in costs and benefits influence the timing and speed of reward-oriented runs, and provided evidence for a complementary contribution of the dorsal and ventral striatum to such adaptive vigor modulations. Before discussing the main results, an important feature of the animals’ behavior must be emphasized, as it constrained how run timing was quantified. In our task, rats produced a series of runs with short inter-run duration (i.e. “bursts” of runs), interleaved with longer pauses. Consequently, the distribution of inter-run durations was highly skewed, with frequent short intra-burst and rare long inter-burst pauses. This pattern of run initiation was well captured by a Wald distribution parameterized by α and γ, which correspond to the decision threshold and drift rate of a drift-diffusion model (DDM), respectively. These parameters should be thought of as descriptors of inter-run duration distributions, analogous to the mean and variance for Gaussian distributions, with the added property that both can determine the peak and spread of inter-run durations (i.e., the pattern of run initiations) in a way that can be intuited by reference to a stochastic accumulation of “motivation” toward a threshold. For example, an overall decrease in run frequency can be accounted for by a concurrent increase in α (higher threshold of motivation to reach to initiate a run) and decrease in γ (slower accumulation of motivation). In contrast, a selective increase in inter-burst pauses with preserved intra-burst frequency can be explained by a decrease in α together with a marked decrease in γ. Within the established framework of utility-based modulation of motor control [3], and given the distinctive running patterns observed in our task, our study yielded four main insights that will be discussed in more detail below. First, across animals, both run frequency and speed declined over the course of a session, in line with the idea that within-session vigor reflects the spontaneously evolving utility to run. Second, animals showed marked inter-individual differences in how they adapted to unsignaled changes in reward delivery probability. While all rats eventually made longer pauses following unrewarded outcomes, “persisters” maintained their intra-burst run frequency and ran faster in low-rewarding blocks, whereas “quitters” reduced their overall vigor. Such idiosyncratic adaptations confirm that, together, run timing and speed (i.e., vigor) reflect the subjective utility to run. Third, all rats spontaneously increased their running speed when motor cost was elevated (e.g., greater platform distance), an adjustment that was not accompanied by a corresponding modulation of run timing. This reveals a fundamental difference in how cost and reward shape behavior: higher motor costs did not discourage running, and the observed increased running speed may reflect an effort to minimize travel time to more distant rewards. Finally, striatal lesions produced complementary vigor deficits: dorsal lesions primarily impaired cost-driven modulation of running speed, whereas ventral lesions blunted the early-session urgency to accumulate rewards, when water is most valuable.

### Rats’ vigor tracks subjective utility during self-paced foraging

In our self-paced foraging task, in which water-restricted rats had to make a series of runs to accumulate drops of water, both inter-run intervals and run durations systematically increased over the course of the session (Fig. 1H, J). This progressive decrease in foraging vigor is straightforwardly explained by a gradual reduction in the utility of running, as animals became increasingly satiated and fatigued. It echoes the well-established observation that vigor reflects the motivational state of the animal [1], with the key difference that, here, it is driven by the animal’s evolving internal state rather than by externally imposed changes in reward rate. This distinction carries important biological implications: when reward availability is low, a thirsty and unfatigued animal may need to sustain a substantial level of behavioral activity in order to maintain reward intake or detect improvements in reward contingencies. While we expect similar patterns to emerge across a wide range of tasks and species, we were surprised to find few, if any, quantitative descriptions of such a parallel modulation of action timing and speed. In humans, this may reflect the common practice of structuring tasks around a fixed number of trials and point-based feedback, which limits the emergence of satiety effects. In other animals, it may stem from a focus on periods of relatively homogeneous motivational states. Interestingly, although both α (run initiation threshold) increased and γ (drift rate toward run initiation) decreased over the session, only changes in α were correlated with the increase in run duration (α_t_ correlated with µ_t_, Supplementary S4D). Our experimental design, in which effort and reward were not perfectly correlated, further allowed us to dissociate their contributions: changes in α were primarily explained by satiety, whereas changes in µ_t_ integrated both effort expenditure and reward accumulation. Overall, these results extend the idea that perceived utility coherently modulates both the timing and the speed of actions to self-paced short locomotor bouts, and incidentally validate our statistical framework for characterizing run timing.

The value of modeling inter-run duration distributions was further illustrated by examining how rats reacted to transitions between high- and low-reward probability blocks. Although all rats ran less frequently during low-reward blocks (positive sawtooth pattern in Fig. 1H, Supplementary S2C), consistent with the known effect of reward rate on vigor [1], the underlying changes in α and γ showed striking and unexpected heterogeneity. On the one hand, all animals showed a reduction in γ under low-rewarding conditions. On the other hand, 9 out of 19 animals displayed a decreased α (α*_u_* < 0). Animals showing the largest decrease in the threshold α exhibited the strongest decrease in the drift rate γ (Supplementary S4A). Since these two parameters have opposing effects on inter-run duration, their coordinated reduction resulted in the smallest net change in overall inter-run duration during low-reward blocks, consistent with longer pauses between bursts whose intra-burst frequency is preserved. Strikingly, this persisters vs. quitters distinction in run timing was mirrored by changes in running speed: persister animals maintained or increased their running speed during lowreward blocks, whereas quitters showed a reduction (Supplementary S4E). The fact that about half of the rats increased their running speed during low-reward blocks may seem at odds with both the normative relationship between reward rate and vigor [1] and the repeated observation that reward-directed movements are executed faster than unrewarded ones [10]. It should be noted, however, that although persister animals ran faster, they also performed fewer runs overall due to the emergence of longer inter-burst pauses, while quitters reduced both the number and speed of reward-oriented runs as expected. A likely explanation for the persisters’ strategy lies in a key difference from previous studies, in which reward availability was explicitly cued. Here, transitions between blocks were uncued, and faced with a series of unrewarded runs, persister animals may have increased their running speed in an attempt to compensate for their lower capture rate, a strategy reminiscent of the frustration effect, whereby unexpected reward omission transiently invigorates subsequent actions [61, 62]. Together, these findings reinforce the idea that running speed provides a sensitive readout of how rats perceive reward utility, which can vary idiosyncratically depending on how unrewarded outcomes are interpreted. Notably, all rats displayed decreased γ following unrewarded runs regardless of their speed strategy, and the magnitude of this effect was uncorrelated with changes in running speed (Supplementary S4F, right), indicating that some aspects of run timing are independent of movement speed. This extends the observation that the speed of decision-making and movement can be coupled or decoupled depending on context [71–75] to freely moving rodents engaged in self-paced foraging.

### Rats flexibly adjust their running speed to maintain reward rate stable despite increasing motor cost

The partial independence between run timing and speed was also illustrated by the observation that changing the distance between platforms or the velocity of the treadmill belt (i.e., the energy and time required to cross the treadmill) had a strong effect on running speed but did not alter the pattern of run initiations (i.e., α and γ remained stable across distances or across imposed belt velocities). Within a utility framework for motor control, the fact that rats ran faster under more effortful conditions might appear counterintuitive, as increased motor cost could have been expected to produce longer inter-run intervals and slower movement (reduced utility to run, [11, 12]). Other studies have nevertheless shown that subjects reaching for distant target distances produce faster movements in order to reduce the cost of time [76, 64]. Our results are congruent with this latter view: increasing the distance to the reward platform seems primarily to be perceived as a potential loss of time, which animals counteracted by running faster. Importantly, although run duration was co-modulated with the threshold for run initiation (α) by the value of crossing the treadmill (µ*_u_* and µ*_t_* correlated with α*_u_* and α*_t_*), variations in running speed brought about by changes in motor cost occurred without correlated changes in α (or γ). This suggests that rats adapt to alterations in perceived reward opportunity and cost through distinct mechanisms. That said, we do not mean to imply that motor cost cannot affect run timing altogether; the constant timing through motor cost manipulation may be explained by the relatively short distances animals had to cross and the spontaneous tendency of rats to save time when faced with more distant targets. Consistent with this, we found that γ (drift rate toward run initiation) decreased with the total effort expended during the session but not with reward accumulation (which primarily accounts for the session-wise increase in running threshold α), confirming a relative partitioning of the mechanisms by which benefit and cost modulate foraging behavior.

It could be argued that the decoupling of run timing and speed observed following motor cost manipulation simply reflects a fundamental difference between locomotion and discrete reaching movements: in reaction time tasks, movement preparation is tightly linked to execution parameters, whereas in our task, the mechanisms controlling speed and those governing run initiation are likely more independent. Moreover, the energetic cost of brief ballistic movements is much smaller than those associated with locomotion. Several observations mitigate this concern: we observed systematic correlations between run timing and speed when the perception of reward changed; the distances covered remained short (less than one meter); and coherent utility-based modulation of speed and decision timing has been reported across motor effectors and species [77]. Moreover, the shape of the speed profile was strikingly conserved across motor cost conditions. Our results should therefore be interpreted not as contradicting findings from other effectors, but as insights afforded by studying locomotion in freely moving animals.

Perhaps our most striking finding is that animals were remarkably flexible in adjusting their speed to maintain a fixed reward rate (e.g., running more slowly when assisted and faster when opposed by the belt). This type of modulation was well-captured by a simple optimal control model in which animals minimize both the cost of time and the cost of effort. Although one might have expected animals to exploit the treadmill assistance and maximize reward by maintaining their speed, this measure instead decreased under this condition. One important experimental consideration is that rats were first trained on the stationary treadmill before being tested with the moving belt; it is therefore possible that when the belt velocity was manipulated, animals attempted to maintain a learned traveling speed acquired during previous sessions, potentially through self-motion cues [78–80]. More generally, the strong modulation of speed by belt velocity suggests the involvement of processes specifically constraining movement speed.

### Complementary contributions of dorsal and ventral striatum to vigor adaptations

The dorsal and ventral striatum (DS, VS) have both been implicated in vigor control [52, 81, 53, 82–84, 2, 3, 85], yet their specific contributions remain difficult to disentangle, partly because vigor deficits can arise from multiple non-mutually exclusive factors (e.g., changes in reward valuation, altered sensitivity to time, impaired perception of motor costs, deficits in motor accuracy or procedural memory). Building on our characterization of foraging dynamics, we exploited the experimental accessibility of rodent models to combine deep behavioral quantification with causal perturbations on a relatively large number of individuals (*n* = 34) with the aim to investigate whether distinct striatal regions contribute to the cost-benefit modulation of vigor. We performed bilateral excitotoxic lesions of either DS or VS. Both lesion groups showed reduced foraging vigor compared to control animals, but with distinct patterns. VS-lesioned rats exhibited markedly longer inter-run intervals and run durations from session onset, both remaining stable throughout, suggesting a reduction in the internal drive promoting frequent and fast running. DS-lesioned rats, by contrast, showed normal early run frequency and speed, with a selective increase in run duration emerging under the highest effort condition. This dissociation was further captured and refined by model-based analyses tracking changes developing within sessions and across cost conditions. Linear discriminant analysis revealed that the three groups could be separated, albeit with some overlap, by a combination of specific parameters. VS-lesioned animals were distinguished by reduced modulation of α and γ with time (both α*_t_* and γ*_t_* close to 0), reflecting diminished urgency to run and modulation over the sessions. DS-lesioned animals were distinguished by increased β, reflecting a reduced tendency to run faster under the most effortful conditions. Finally, both lesioned groups showed increased ς_0_, indicating a reduced willingness to incur effort costs to save time, an effect more pronounced in VS-lesioned animals, consistent with an overall reduction of reward urgency. These results bear on longstanding assumptions about DS versus VS contributions to reward-oriented behavior, and on the type of vigor deficits observed in Parkinson’s disease, which are believed to primarily reflect DS dysfunction.

The overall reduction in urgency to drink at session onset following VS lesions is in line with the well-established contribution of the VS to appetitive and motivational processes such as approach behavior, sustained pursuit of reward, and sustained work [49, 81]. Specifically, dopamine in the VS is necessary when animals must sustain work to obtain rewards in progressive ratio schedules [25], though in such tasks, motor cost is minimal and whether deficits reflect impaired effort production or decreased urgency is difficult to disentangle. Consistent with our observations, alteration of VS projection neuron activity prolongs the time to initiate and complete lever-press sequences [86], paralleling the longer inter-run and run durations and the pronounced effort-time tradeoff (ς_0_) we report. Altogether, and given that VS-lesioned animals showed no specific deficit in increasing their speed under effortful conditions, these results argue for an effect of VS lesions on time sensitivity (urgency) rather than on effort per se. Strikingly, VS-lesioned animals behaved at session onset as if satiated or fatigued. Consistent with this, dopamine in the VS core signals reward-predicting cues selectively in non-satiated animals [87], and D1/D2 SPNs in this region are strongly activated in food- and water-restricted animals given the opportunity to consume rewards [86]. Our results therefore support the view that the VS integrates environmental information with signals reflecting internal state to adapt the willingness to work accordingly [88, 49, 89]. One advantage of our rich cost-benefit framework is that it also exposes preserved capacities in lesioned animals: VS-lesioned rats still modulated their vigor according to reward availability, suggesting this sensitivity relies on other circuits. Finally, VS lesions produced a marked increase in run duration variability during low-reward blocks. Given that movement variability increases following unrewarded outcomes [59], this result suggests that the VS plays a key role in constraining the speed of reward-oriented movements as a function of the discrepancy between internal state and reward opportunity.

Turning to rats with DS lesions, their primary behavioral difference from both control and VS-lesioned animals was a limited increase in running speed under the most effortful conditions, captured by increased β values reflecting greater sensitivity to motor cost. This is particularly relevant to bradykinesia and akinesia in PD, which has been proposed to reflect an adaptive response to increased effort sensitivity [18, 19, 90], assuming that striatal lesions increase basal ganglia output activity. One might ask whether this pattern instead reflects a primary locomotor impairment, given the generally assumed motor functions of the DS (e.g., [91–93]). Several observations argue against this. A general locomotor deficit would be expected to affect all displacements irrespective of velocity, which is not what we observe. Extensive lesions of the dorsolateral striatum do not significantly alter locomotion assessed with 3D tracking [94], consistent with a modulatory rather than generative role of basal ganglia in locomotion [95]. DS-lesioned rats retained the ability to adapt their locomotion on a treadmill moving at increasingly faster speeds [35]. In our task they displayed a preserved running speed profile and could still reach high running speeds in our task, arguing against a locomotor impairment preventing them to run quickly or changing qualitatively how they run. Crucially, β values of control and DS-lesioned animals showed substantial overlap: some non-lesioned control animals exhibited high β values comparable to DS-lesioned rats, making it unclear why similar behavioral profiles should be attributed to motivation in one case and locomotor deficit in the other, an argument that applies equally to VS-lesioned animals (i.e., why reduced vigor in VS-lesioned animals should not be interpreted as a locomotor deficit). Our cost-minimization model thus provides a parsimonious framework to interpret both inter-individual and inter-group differences without invoking ad hoc explanations. It is also worth noting that the interpretation of bradykinesia as increased effort sensitivity has been challenged [21], with the alternative proposal that PD motor deficits arise from impaired transitions between stable and dynamic movement states. Because such transitions require a cost, this interpretation remains compatible with the DS being critical to adjust vigor according to motor cost. Regardless of the interpretive framework, the observation that DS-lesioned animals display reduced scaling of their running speed according to cost conditions aligns with the function of the anterior cingulate cortex in integrating effort [96], which strongly projects onto the dorsal striatum [97] and with human studies implicating the DS in encoding the cost of effortful actions [20, 98]. While our results draw parallels with bradykinesia, Parkinsonian motor symptoms also include episodic pausing and inability to sustain continuous locomotion (festination and freezing of gait). DS-lesioned animals did not display overt locomotor arrest during runs, even under the most effortful conditions.

However, animals with larger DS lesion displayed higher baseline values of both ς and α (Supplementary S7I, J), suggesting that larger DS damage jointly affects run timing and speed, more closely resembling the combined akinesia and bradykinesia characteristic of Parkinsonism. More broadly, given that distinct striatal regions have been implicated in a wide range of motor and motivational disorders, from Parkinson’s disease to depression and addiction [2], combining naturalistic foraging paradigms in rodents with cost minimization modeling offers a computationally tractable framework to understand how basal ganglia neuronal types and pathways shape decisions and movements, with strong translational potential [99, 100].

## Methods

### Animals

A total of 53 wild-type Long-Evans rats (Janvier Labs, France) were used (males *n* = 29, females *n* = 24). They were ∼12 weeks old at the beginning of the experiments and housed in groups of 2–3 rats in temperature-controlled ventilated racks. A reversed 12 h–12 h light/dark cycle was maintained and all experiments were carried out during the dark phase. Food was available ad libitum in their home cage. The animals underwent a controlled water restriction protocol once engaged in the foraging task (see Methods for details). Body weight was measured twice a day. During water restriction periods, the rats consistently maintained their body weight at, or above, 85% of their pre-restriction weight. Experiments were carried out according to standard ethical guidelines for animal experiments (European Community Directive 86/609/EEC) and were approved by the national ethics committee (Ministère de l’enseignement supérieur et de la recherche, France, Authorization # 16195 and # 46285).

### Task Apparatus

Two corridors placed in sound-attenuating boxes were used for the experiments. They were 120 cm long and 14 cm wide, surrounded by 70 cm high plexiglass walls. The floor of these corridors was in fact the belt of a treadmill driven by a brushless digital motor (BRG 44 SI, Dunkermotoren). The belt was flanked by two platforms (15 cm long) on which the rats could rest and/or drink (Fig. 1A). Two reward ports affixed to the front and rear walls were connected to solenoid valves (VAC-20 PSIG, Parker) and dispensed calibrated drops of water (12 µL). One of the walls and platforms could be moved to modify the distance between the two reward ports. A strip of LED lights installed on the ceiling along the treadmill provided lighting. The rat’s position was tracked by a ceiling-mounted camera (DMK 23GM021, The Imaging Source, 25 fps) and was defined as the centroid of its body. Reward zones were defined by virtual boundaries placed 20 cm from each end of the corridor (5 cm before the platforms). Crossing a boundary triggered water delivery and, in the treadmill-on condition, halted for a second and reversed the treadmill, ensuring that belt motion was always consistent relative to the rat’s running direction. This pause allowed rats to safely step onto the platform without risk of injury. The entire apparatus and data recording was fully automated using a custom-made program (LabVIEW, National Instruments).

### Foraging experiments

Each rat was handled ∼30 minutes each day for 3–5 days before familiarization with the foraging task. The water bottles were replaced with water with 0.5% citric acid at the beginning of handling and completely removed at the beginning of testing [101]. Rats were habituated to the task rules twice a day for 2 days, in 30 min sessions. During these sessions, the distance between the reward ports was 90 cm and the treadmill belt remained immobile. The rats had to alternate between the two reward ports to receive a single drop of water (∼12 µL) after each crossing (Fig. 1A). The rats understood this rule very quickly as all of them already ran back and forth between the two reward ports and consumed the drops of water at the end of the second, if not first, habituation session. During the main behavioral task, water-restricted rats had to run back and forth between the two reward ports to receive drops of water. They performed the task twice a day in 1 h sessions. Within a session, the probability of receiving the reward after each crossing of the apparatus alternated between high (90%) and low (10%) in 5 minutes-long uncued blocks.

Every day, the effort (action cost) required to obtain each reward was manipulated. In a first set of experiments, the distance between the reward ports was set to either 60, 90 or 120 cm, corresponding to a physical distance run of respectively 29, 62 and 94 cm. Each distance was repeated for 3 days (that is, 6 sessions in total as rats performed two sessions per day). The order of the distance conditions was pseudo-randomized but kept constant across rat batches. The range of physical distances that can be tested in our apparatus is limited and we used the motorized treadmill belt to manipulate motor cost without changing travel distance (Methods). After a short break (2 days with full access to food and water), the rats were familiarized with crossing the corridor while the treadmill belt was turned on. First, the treadmill was activated very slowly in the same direction as the running direction. The speed of the treadmill belt was gradually increased during training to avoid stressing the animals and to prevent injuries. Then, animals were familiarized with running while the belt direction counteracted their movements, following the same protocol. This familiarization phase lasted for 4–6 days in 1 hour-long sessions, 2 sessions per day. Animals were water-restricted and the structure of the session followed the high/low probability block described above. Following this familiarization phase, in a second set of experiments, rats performed two sessions per day, during which the treadmill belt was activated at a fixed velocity relative to the animal’s running direction (20, 10, ±2, −10 or −20 cm/s). That is, the treadmill direction alternated each time the animals reached a platform. Every day a new belt velocity was used, each treadmill velocity was repeated for 3 days (6 sessions in total for each velocity). In pilot experiments we found that the foraging behavior of animals was indistinguishable at the two lowest speeds (±2 cm/s, the similarity was confirmed in the animals tested in this study), and those conditions were therefore pooled together. Rats performed 6 sessions in total in those two lowest speeds (typically 3 sessions at 2 and 3 sessions at −2 cm/s). The order of the conditions was pseudo-randomized but kept constant across rat batches. 19 rats (males *n* = 11, teal gradient; females *n* = 8, violet gradient) were used as control with no removal of any individuals from the study. Each rat performed *n* = 48 sessions in total (6 sessions × 8 action cost conditions).

### Data processing

Corrupted sessions (e.g., poor video tracking due to urine reflection or computer system crash, *n* = 105 sessions, 4.12% of all sessions) were excluded from the analysis. The position of the rat was smoothed with a Gaussian kernel (σ = 2 samples). The instantaneous speed of the rat was computed as the distance traveled between two successive frames, divided by the time between two successive frames (i.e., 0.04 seconds). The resulting speed profiles were subsequently smoothed with the same Gaussian kernel (σ = 2 samples) to reduce high-frequency noise arising from discrete sampling; empirically, this two-step procedure did not erase the main features of the velocity profiles and simplified downstream processing. In the “treadmill on” conditions, the running speed of the rat was obtained by subtracting the velocity from the treadmill belt to the animal’s speed in the fixed reference of the experimental room.

The inter-run and run periods were separated using a double threshold based on the instantaneous speed and position along the apparatus. For each session, we computed a two-dimensional kernel density estimate (KDE) of speed as a function of position. This revealed high-density regions corresponding to low-speed epochs near the extremities of the apparatus. We then applied a fixed density threshold to extract these regions and, for each position, derived a corresponding speed boundary. These position-dependent boundaries defined inter-run zones on both sides of the track. Time points falling within these zones were classified as inter-run periods, whereas all remaining epochs were classified as run periods. Brief spurious transitions between run and inter-run states were subsequently corrected based on spatial continuity. The average running speed of the rats when crossing the corridor (e.g., in Fig. 2F) is defined as the distance traveled during a run period divided by the run duration. The median speed profile computation was implemented from the Karasec algorithm found in [102] (Fig. 3).

In each plot, individual data (colored lines, color code per animal maintained throughout the paper), group median (thick lines) and interquartile range (error bars) are represented. The number of rats in each group is indicated in the figure or in the figure legends. Statistical analyses were performed with custom Python scripts using permutation tests (in every case *n* = 10000 iterations).

### Inter-run duration model

Related to Fig. 1. For each rat, we wanted to quantify the dynamics of inter-run durations throughout the session, its modulation by the probability of receiving a reward when crossing the treadmill, and by the action cost conditions (3 distances between the platforms and 5 treadmill belt velocities). An example inter-run duration distribution from the first 10min of the example session is shown in Fig. 1C. We fitted this distribution with a Wald distribution [103] (black). The choice of a Wald distribution was driven by the fact that, while median values can show broad trends, they fail to reflect changes in the shape of the underlying distribution, and by the observation that it provided better fits than candidate alternatives including log-normal and Lévy distributions. The α and γ parameters of the function correspond to, respectively, the threshold and rate of accumulation of a one-bound drift-diffusion model (DDM) [104, 56, 57] capturing the decision process until run initiation (i.e., inter-run duration). Indeed, the explicit solution resulting from the diffusion process is the Wald distribution [103] with probability density function:

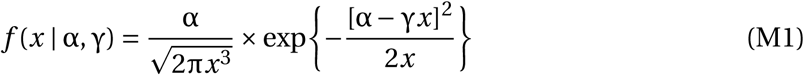

The one-bound DDM describes a continuous time-stochastic accumulation process in which a quantity *x* (in our case motivation) continuously accumulates until it reaches a threshold α. *x* is accumulated with a drift rate γ with Gaussian noise. Once *x* reaches the threshold α, the animal initiates a run. This process is simulated (*n* = 50 simulations) in Fig. 1C, inset. Although other types of distributions, such as the log-normal distribution, provided qualitatively similar descriptions of the inter-run durations, we chose the Wald distribution as it provided an intuitive interpretation of the decision process governing the inter-run duration patterns observed in our experiments.

For each rat and each experimental condition, the initial values of α and γ may be different. We named α_0_ and γ_0_ the values of α and γ at the beginning of the session (Fig. 1E).

Furthermore, α and γ can increase/decrease throughout the session (e.g., rats may need to accumulate more motivation at the end of the session when they are satiated). To assess the effect of time *t*, the complete inter-run duration distribution is further separated into 6 bins of 10 minutes. This allowed for a good resolution for the temporal evolution in the session while ensuring that both high and low reward probability blocks are represented in each time bin. This evolution is captured by α_t_ and γ_t_ (Fig. 1E).

Finally, α and γ can be weakly or strongly modulated by the recent reward outcome history (e.g., rats may accumulate motivation slower when the probability of receiving a reward decreases from 90% to 10%). Therefore, each time bin was further separated according to the history of unrewarded runs, *u*, defined as the ratio of runs that were not rewarded over the 3 previous runs (Fig. 1B, top); it ranges from 0 (the rat received 3 rewards out of its previous 3 runs) to 1 (the rat received no reward out of its 3 previous runs). The choice to consider 3 previous runs is justified by the minimization of the error of the inter-run duration model (Supplementary S1C). We named α_u_ and γ_u_ the magnitude of the change in α and γ when the number of unrewarded trials increased (Fig. 1E).

We defined the following model in which α and γ capture the complete inter-run duration distribution at time bin *t* and history of unrewarded runs *u*:

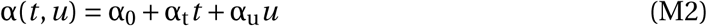

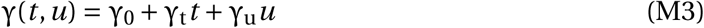

For each rat in each experimental condition, we computed α_0_, α_t_, α_u_, γ_0_, γ_t_, and γ_u_ that mini-mized the total error of the model using maximum likelihood. The minimization procedure was achieved using the L-BFGS-B algorithm implemented in the SciPy module [105], with the constraint that α_0_ and γ_0_ are positive.

### Run duration model

Related to Fig. 1D. The run duration model follows a procedure similar to that described for the inter-run duration model. Run duration distributions were fitted with a Cauchy distribution with median µ and half-width at half-maximum σ (i.e., the variability of the run durations). The Cauchy distribution is similar to a Gaussian distribution but with a higher peak and heavier tails, which allowed us to capture rare long run durations, and provided a better fit than Fréchet or ex-Gaussian distributions. Although the experimental distributions show some left asymmetry reflecting the minimum time physically required to cross the apparatus, this constraint is identical across all animals within a given condition and does not introduce systematic bias in parameter estimations or group comparisons.

As for the inter-run model, the parameters describing the run durations of each rat and each experimental condition can have a varying initial value of µ and σ (µ_0_, σ_0_), they can increase/decrease throughout the session (µ_t_, σ_t_), and may be modulated following a series of unrewarded runs (µ_u_, σ_u_).

We defined the following model in which µ and σ captured the complete run duration distribution at time bin *t* and history of unrewarded runs *u*:

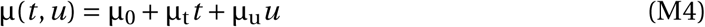

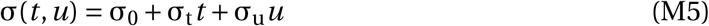

We computed the best µ_0_, µ_t_, µ_u_, σ_0_, σ_t_ and σ_u_ that maximized the likelihood of observing the experimental run duration distributions in each experimental condition (Supplementary S1).

### Quantification of model goodness of fit

Related to Supplementary S1. Model fit was computed by taking the median of synthetic observations generated using the fitted parameters. This procedure was repeated to obtain a confidence interval (*n* = 10000 iterations); in each iteration, the number of synthetic observations generated in each block was equal to the sum of experimental data points aggregated over the 6 sessions of the 94 cm distance condition (Supplementary S1A, S1E). We estimated the error (in seconds) between model fit and experimental data for the inter-run and run duration models. The absolute difference between the observed experimental median and the resampled medians was computed in each block for all rats (*n* = 19, Supplementary S1B, S1F). The median error is around 0.6 seconds for the inter-run duration model and 0.04 seconds for the run duration model. To define the optimal number of trials in memory, we aggregated data from all experimental conditions and separated them according to whether previous runs were rewarded or unrewarded. This procedure was performed with trial history lengths ranging from 1 to 8 and the Bayesian Information Criterion (BIC) was computed for each rat and each history length condition in both the inter-run and run duration models. We chose to consider the 3 previous runs as this is the number that minimizes the error of the inter-run duration model (Supplementary S1C, S1G). For both the inter-run and run duration models, a likelihood ratio test was used to determine if removing the parameters α_t_, α_u_, γ_t_, γ_u_, µ_t_, µ_u_, σ_t_ and σ_u_ significantly decreased the accuracy of the models. Experimental data from all animals in each experimental condition were pooled to compute the log-likelihood for the complete models (with all parameters free) and a set of nested models (with one of the parameters fixed to 0). The log-likelihood ratio between the complete model and each nested model was computed as

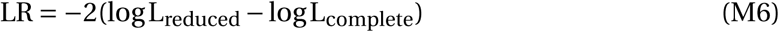

and statistical difference in goodness of fit between the complete model and each of its nested models was assessed using a χ^2^ distribution with degrees of freedom *df* equal to the difference in the number of parameters between these two models (Supplementary S1D, S1H).

### Model-free analysis

Related to Supplementary S2. We performed a complementary model-free analysis to assess whether the effects captured by the model-based approach were directly observable in the data. For each animal, inter-run and run durations distributions were pooled across all sessions of a single experimental condition (120 cm, treadmill off). Model-free parameters were computed by fitting Wald distributions to inter-run durations and Cauchy distributions to run durations. For time-dependent parameters (α_t_, γ_t_, µ_t_, σ_t_), data were pooled into 10 min bins. Distribution parameters were estimated independently for each bin, and the corresponding model-free parameter was defined as the average change in the estimated parameter between consecutive bins. For reward-dependent parameters (α_u_, γ_u_, µ_u_, σ_u_), data were separated into low- and high-reward probability blocks. Distribution parameters were estimated separately for each reward condition, and the model-free parameter was defined as the difference between parameters obtained in low- versus high-reward probability blocks.

Model-free parameters were then correlated with their corresponding model-based parameters across animals. Statistical significance was assessed using a permutation test (*n* = 10000), with two-sided *p*-values computed from the distribution of permuted correlation coefficients.

### Model simulations

Related to Fig. 1H, J. As the experimental data was fitted in 6 time bins of 10 minutes (thus containing a block of high and a block of low reward probability), we extrapolated the values of α(*t*, *u*) and γ(*t*, *u*) in each 5 min block, by assuming that high probability blocks mostly contained *u* = 0 and that low probability blocks contained *u* = 3.

We then used the α(*t*, *u*) and γ(*t*, *u*) functions to draw synthetic inter-run durations (same number of observation as the experimental data) from the Wald distribution (*n* = 10000 resamplings, Fig. 1H). The same method was used with the fitted µ_0_, µ_t_, µ_u_, σ_0_, σ_t_ and σ_u_ parameters of the µ(*t*, *u*) and σ(*t*, *u*) functions to draw synthetic run durations from the Cauchy distribution (Fig. 1J).

### Inter-block inter-run modulation: model-free assessment

Related to Supplementary S4. To correlate the parameter α_u_ with the modulation of inter-run duration between reward probability blocks, a model-free estimate of this modulation was computed. For each animal, data from each session performed under the condition where the treadmill belt was immobile and the corridor distance was set to 94 cm were pooled into 10 minutes bins. For each bin, a delta between the median run duration in the low reward probability block (10%) and the high reward probability block was calculated. These pairwise deltas were then averaged using the median across all block pairs within a session, and the resulting per-session values were further collapsed to a single per-animal estimate by taking the median across sessions. Statistical significance was assessed using a permutation test (*n* = 10000), with two-sided *p*-values computed from the distribution of permuted correlation coefficients.

### Parameter correlations

Related to Supplementary S4. We used linear mixed models to compute pairwise correlations between the fitted parameters of the run and inter-run duration models (statsmodels module [106]; Supplementary S4). The conditional *r* ^2^ values were computed using the method described in [107]. The correlation matrix and dendrograms were generated using the Scikit-learn module ([108], Supplementary S4).

### Total estimated effort and water drops collected

Related to Supplementary S5. The total estimated effort and the total amount of water obtained in each experimental condition were computed for all rats (*n* = 19). The total amount of water obtained was defined as the sum of water drops collected in a session. The total effort expenditure for each session was estimated as the sum of the square of the instantaneous speed *v* (*t*) [65] of the rat at each frame *f* of the session (sessions were 1 hour long and thus contained 90000 frames), multiplied by the average weight *w* of the rat:

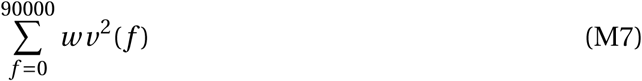

The median total effort (Supplementary S5A) and total water (Supplementary S5B) of all sessions across action costs is represented for each rat.

### Correlation between total estimated effort and water drops collected and behavioral parameters

Related to Supplementary S5. To determine whether modulations in time-dependent behavioral parameters (α_t_, γ_t_, µ_t_, σ_t_) were better explained by total effort expended (E_TOT_) or total rewards obtained (R_TOT_), we fitted linear mixed-effects models using maximum likelihood estimation (restricted maximum likelihood disabled; [106]). For each variable, the full model included both predictors, with animal included as a random intercept:

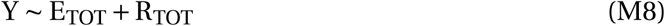

To assess the unique contribution of each predictor while controlling for the other, we computed the fixed-effect coefficient β, its 95 % confidence interval and associated *p*-value from the full model. For visualisation, model predictions were generated while holding the nontested predictor at its mean value. Confidence bands for predicted correlation were obtained by resampling (10000 iterations) animals with replacement and refitting the model at each iteration.

Model comparison was performed using Akaike Information Criterion (AIC). We compared the full model against reduced models containing only (E_TOT_) or only (R_TOT_). Differences in AIC (ΔAIC) were computed relative to the full model and interpreted following Burnham and Anderson ([109]), with ΔAIC thresholds of 2 (weak), 4 (moderate), and 7 (strong evidence).

### Costs minimization model of run durations

Related to Fig. 3. The speed profile *v* (*t*) of the rats has the same shape in all experimental conditions and can thus be described with a universal speed profile (Fig. 3B). The universal speed profile *ṽ*(*t̃*) is scaled in time by the run duration T and by the maximum velocity V_max_, such that:

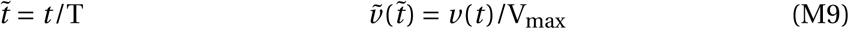

At each instant *t*, the rat travels on the treadmill belt (which moves at speed *v*_belt_) with a velocity *v* (*t*) = *v*_belt_+V_max_*ṽ*(*t̃*) in the laboratory framework, to reach the final position *x*(T) = *d*. By integrating *v* (*t*), the maximum velocity attained during this movement can be expressed as a function of T, *v*_belt_ and *d* as:

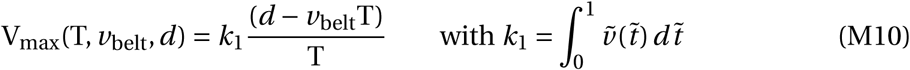

The parameter *k*_1_ only depends on the universal speed profile *ṽ*(*t̃*) and is thus a constant. We defined an energetic cost of movement C_M_ (Fig. 3C, red curve) that is proportional to the relative velocity of the rat (*v* − *v*_belt_) raised to some power β (Fig. 3D). The amount of energy spent walking on a flat surface has been shown to correspond to a fixed exponent value of the running speed (i.e., β = 2; [65]), but we chose to leave this exponent free. An increase in β increases the cost of fast velocities. C_M_ is expressed as:

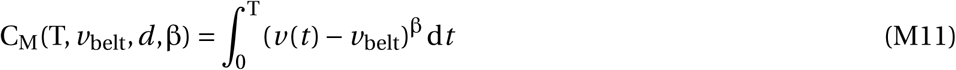

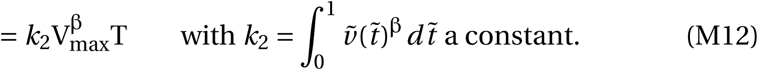

Note that *k*_1_ and *k*_2_ depend only on the shape of the universal speed profile *ṽ*(*t̃*) and are therefore constants; the exact shape of the speed profile does not affect model predictions, as it only contributes fixed scaling factors to V_max_ and C_M_. This parametrization yields two simple and interpretable parameters that can be estimated from individual animals and shares the same mathematical structure as classical metabolic cost models for quadrupedal mammals ([67]) without requiring precise estimates of joules or oxygen volume consumption. Although energetic cost is known to scale with body mass ([110, 111]), mass was not incorporated as an explicit parameter to keep the model parsimonious. Post hoc analysis confirmed that mass-normalizing ς (i.e., 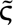 = ς/*w*) does not alter the group differences between intact, DS-, and VS-lesioned animals, ruling out body mass as a confound in our lesion comparisons. The sensitivity of parameters to body mass, which itself covaries with sex in rodents, opens an interesting perspective for future studies on how physical characteristics modulate inter-individual variability in the motivation to initiate movement. We defined a second cost related to the passage of time C_T_ that increased monotonically with the duration of the movement T (Fig. 3C, green curve). We chose:

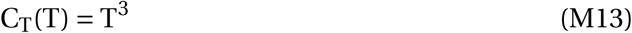

This formulation was selected because it yields a closed-form analytic solution for the optimal run duration T_opt_; alternative formulations such as T^2^ or T^−1^ do not cancel cleanly with C_M_and preclude an analytic optimum. This choice is moreover consistent with established cost minimization frameworks in motor control ([9]). The exponent (here, 3) could in principle be treated as a free parameter, but this would require numerical optimization and sacrifice interpretability. In a running epoch, we assumed that the rat minimized the total cost C, defined as the sum of the cost of movement C_M_ and cost of time C_T_ (Fig. 3C, black curve):

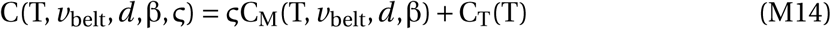

The parameter ς is the relative sensitivity between the costs of movement and time (Fig. 3D). Note that an increase in ς is mathematically equivalent to a decrease in the relative weight of C_T_, and the model cannot distinguish between these two interpretations from run duration data alone. The total cost C is a concave function of T, meaning that there is an optimal running duration T_opt_ that minimizes the sum of the two costs (Fig. 3C). T_opt_ can be expressed as a function of distance *d*, treadmill belt *v*_belt_, and the relative sensitivity to motor cost ς:

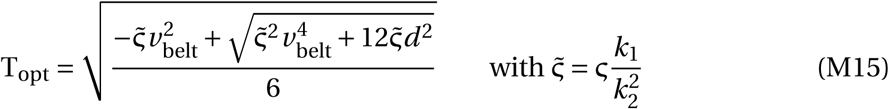

By varying the parameters *d* and *v*_belt_, we were able to predict this optimal crossing duration for a given ς and β.

For each animal, we computed the parameters ς_0_ (initial motor-cost sensitivity), ς_t_ (evolution of motor-cost sensitivity throughout the session), ς_u_ (change of motor-cost sensitivity with reward outcome history), and β that minimized the sum of squared error between experimental µ_0_, µ_t_, and µ_u_ for each condition and the prediction generated by this model. The minimization process was achieved using the SciPy minimize module [105] and the Nelder-Mead algorithm. For each rat, we chose to fit a single value of β.

### Quantification of effort model goodness of fit

Related to Supplementary S6. To evaluate the accuracy of the effort model across experimental conditions, we analyzed the residual error between model prediction and experimental data. For each animal, residuals were computed using the best-fit parameters of the complete effort model, and defined as the absolute difference between predicted and observed run durations across all conditions. Residuals were averaged across time bins and reward history conditions, yielding for each animal a mean residual error in seconds for each experimental condition.

To assess the contribution of parameters to the effort model, we used likelihood-ratio tests to determine if removing the parameters ς_t_, ς_u_, or fixing β to its classical value (fixed to 2; [65]) significantly decreased the accuracy of the models. Model loss was computed by minimizing the sum of squared errors (SSE, the minimization process was achieved using the SciPy minimize module [105] and the Nelder-Mead algorithm) between model prediction and experimental data across all experimental conditions (N = 8 conditions ×6 time bins ×4 unrewarded history). To compare the nested models to the complete model, SSE were summed across animals. Assuming independent Gaussian residuals, we computed a likelihood-ratio for each nested model relative to the complete model as

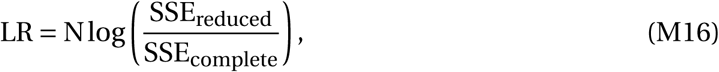

and a statistical difference in goodness of fit between the complete model and each of its nested models was assessed using a χ^2^ distribution with degrees of freedom *df* equal to the difference in the number of parameters between these two models.

### Lesion surgeries

Related to Fig. 4. Excitotoxic lesions of the dorsal and ventral striatum (DS and VS respectively) were performed under deep anesthesia, induced with an intraperitoneal injection of a mixture of 100 mg/kg Ketamine and 10 mg/kg Xylazine and maintained with inhalant isoflurane gas (less than 0.5%). After shaving and cleaning the scalp, the animal was placed in a stereotaxic frame (Kopf instruments). A local analgesic (lidocaine) was injected under the scalp and an incision was made along the midline of the skull. The exposed skull was cleaned and craniotomies were drilled above the target areas.

To perform a bilateral fiber-sparing lesion, ibotenic acid (1% in 0.1 M NaOH, Fisher Scientific) was infused via a thin glass pipette connected to a nanoinjector (Nanoliter2020 Injector, World Precision Instruments) at a rate of 90 nL/min. The stereotaxic coordinates for the injection sites for DS and VS lesions are shown in Table 1. For dorsal injections, the glass pipette was lowered to the lowest of the two dorsoventral levels and a first injection of half the total volume was injected. The pipette remained there for 3 minutes before being retracted 100 µm, and the rest of the volume was infused. After injecting the final volume for each site, the pipette remained in place for 7 minutes before being slowly retracted to prevent drug backflow. Once all injections were completed, the craniotomies were sealed with bone wax, the skull was disinfected, the skin was sutured and painkillers were administered.

**Table 1:** Injection coordinates and injection volumes for DS and VS lesions. Coordinates are measured from Bregma.

|  | DS |  |  | VS |  |
| --- | --- | --- | --- | --- | --- |
| AP | +1.56 | +0.6 | −0.36 | +2 | +1.3 |
| ML | $\pm 2.7\text{--}\pm 3.4$ | $\pm 3.1\text{--}\pm 3.9$ | $\pm 3.5\text{--}\pm 4.1$ | $\pm 1.5$ | $\pm 1.2$ |
| DV | −5.5 & −4.5 | −5.5 & −4.5 | −5.5 & −4.5 | −7.2 | −7.4 |
| Dose | 250–300 nL | 250–300 nL | 250–300 nL | 100–120 nL | 100–120 nL |

The rats were housed individually to prevent possible harm from cagemates for 5 days and were allowed to recover for at least 10 days before the start of behavioral procedures. Lesioned rats performed the same training procedure and experimental sessions as control rats (6 sessions × 8 action cost conditions). In total, 20 rats underwent an excitotoxic lesion of the DS (males *n* = 11, females *n* = 9; green gradient). One lesioned male rat was removed after a couple of familiarization sessions due to a hind leg malformation (unnoticed during homecage handling) and difficulties in running. 15 rats underwent an excitotoxic lesion of the VS (males *n* = 8, females *n* = 7; red gradient).

Non-lesioned rats were used as controls instead of sham-operated animals for two main reasons. First, sham procedures can themselves induce behavioral changes, complicating interpretation [112]. Second, the observation that dorsal and ventral striatum lesions produced qualitatively distinct behavioral deficits provides strong internal evidence that the effects are attributable to the specific region targeted rather than to the injection procedure itself. Together, these considerations justified the use of non-lesioned controls, an approach that also aligns with the 3Rs framework by reducing the number of animals used and limiting unnecessary surgical procedures (Refinement).

### Immunohistochemistry

Related to Fig. 4. At the end of the experiments, the rats were euthanized with an intraperitoneal injection of a mixture of 0.4 mg/kg Domitor and 40 mg/kg Zoletil. They were perfused with 4% paraformaldehyde and their brains were collected for histological analysis of the lesion size and location. Brains were coronally sliced (60 µm thickness) on a vibratome. For each animal, sections corresponding to the location of each injection on the rostrocaudal axis were selected and submerged in 0.1 M PBS.

PBS was replaced with a citrate buffer (10 mM) for 10 minutes at room temperature and the slices were submerged in a blocking solution (5% of normal goat serum into PBS with 0.3% Triton) for 120 minutes at room temperature. The blocking solution was then replaced with another consisting 1/500 of mouse anti-NeuN antibody (Merck Millipore, MAB377) and 1/500 of rabbit anti-GFAP antibody (Agilent, Z033429-2) diluted in 250 µL of the blocking solution per well, and kept overnight at 4 °C. The sections were then rinsed twice for 10 min in PBS-Triton at room temperature before being submerged again in 1/500 of donkey anti-mouse antibody (Al555, red), 1/500 of donkey anti-rabbit antibody (Al488, green) diluted in 250 µL of PBS for 120 minutes at room temperature. Finally, they were washed twice in PBS, for 10 minutes each time, and mounted for microscopy.

Fixed brain slices were imaged using an Apotome microscope (Apotome Imager Z2, Zeiss) and stitched in the processing software (Zen, Zeiss). For each slice, the striatum and lesion area were manually outlined in both hemispheres (Fiji, [113]). The area and centroid of each region of interest were computed. The whole procedure was performed blindly to the behavioral results.

### Max speed distribution comparison

Related to Supplementary S7. To compare the distributions of peak running speeds between the DS-lesioned and control groups (Supplementary S7E,F), we used a non-parametric permutation approach based on kernel density estimation (KDE). For each animal, we pooled all run data and estimated the KDE of peak running speeds using a Gaussian kernel (light curves). To compare difference between groups, we pooled all data across animals within each group; the observed difference in pooled KDEs (DS-lesion minus controls) was compared against a null distribution obtained by randomly permuting group labels across animals (*n* = 10000 permutations). A global confidence band was derived by iteratively tightening the percentile bounds until the proportion of permuted differences exceeding the band fell below 5 %, providing a global 0.05 threshold that controls for multiple comparisons across the speed axis. Portions of the observed difference curve exceeding this global confidence band are considered statistically significant and are displayed in black in the figures.

### Linear Discriminant Analysis

Related to Fig. 4.To assess multivariate group differences across behavioral parameters, we performed linear discriminant analysis (LDA) using group identity (control, dorsal striatum lesioned, ventral striatum lesioned) as class labels. Individual animals were projected onto the first two linear discriminant (LD) axes, which explain the largest proportion of between-group variance. Group centroids were computed as the mean LD scores within each group. Within-group dispersion was visualized using two standard-deviation ellipses derived from the covariance of LD1 and LD2 scores for each group. To visualize class separation, a multinomial logistic regression classifier was trained on the LDA projections, and decision boundaries were computed in the LD space. Classification accuracy was estimated from the classifier predictions. Statistical significance of the classification was assessed using a permutation test in which group labels were randomly shuffled and the classifier accuracy recomputed; the observed accuracy was compared against this null distribution to determine whether it could be observed by chance. Behavioral parameter contributions to the discriminant axes were quantified using the LDA loadings. Loadings for LD1 and LD2 were displayed as vectors, with vector length reflecting the magnitude of each variable’s contributions; variables with high contributions (arbitrary threshold set at 0.6) were displayed in black and the next three more relevant variables are shown in gray.

### Statistical analyses

We performed linear regression with permutation to test whether manipulating effort conditions (3 distances and 5 treadmill velocities) had an effect on each model parameter. Briefly, experimental condition labels were randomly shuffled to estimate the likelihood of obtaining the observed slope and intercept; *p*-value for the slope was defined as the fraction of shuffles with a slope magnitude greater than the observed slope, (Fig. 2).

To compare the medians of two independent groups (i.e., control and lesioned rats), a permutation test was performed by permuting the group labels and computing the probability of obtaining a median difference at least as large as the observed one (Fig. 4D). For paired comparisons (e.g., early vs. late parameters, Supplementary S3), condition labels were permuted within pairs, and the probability of obtaining a median difference at least as large as the observed one was similarly computed. In both cases, the *p*-value was defined as the fraction of permutations yielding a median difference greater than or equal to the observed difference.

When indicated, we applied the Bonferroni correction to mitigate potential Type I errors associated with multiple comparisons; corrected *p*-values are reported (i.e., raw *p*-values multiplied by the number of comparisons) rather than adjusting the significance threshold. *p* > 0.05 was considered not statistically significant. Significant *p*-values are represented in black and non-significant *p*-values in gray.

## Data and code availability

Data from each behavioral session were saved in text files containing the position of the rat and all task parameters. The data processing pipeline was implemented in custom-made Python scripts. Experimental data (text files), hand-drawn illustrations (Fig. 1A, 2B) and notebooks used to process, quantify, visualize behavior, and generate the figures of this manuscript will be publicly available upon publication of the manuscript.

## Acknowledgements

This project has received funding from the French National Research Agency (ANR-22-CE37-0010-01 BasalCost; DR), from the “Investissements d’Avenir” French Government program managed by the French National Research Agency (ANR-16-CONV-0001), from Excellence Initiative of Aix-Marseille University - A-MIDEX (TM) and from the Fondation pour la Recherche Médicale (FDT202204014828; TM). This work was supported by the INMED core facilities Histology and InMAGIC, as well as the CENTURI Multi-Engineering Platform. We thank Marie Kurtz for brain slicing and immunohistochemistry procedures, and Élodie Fino, Ambre Linosier and Julie Buron for technical help with intracerebral injections. We also thank Kenza Amroune, Jérôme Epsztein, Lorenzo Fontolan, Lionel Rigoux, Mostafa Safaie, and Maud Schaffhauser for critical reading of this and earlier versions of the manuscript, and the Cortico-Basal Ganglia Circuits and Behavior team for discussions.

## Author contributions

**Thomas Morvan:** Conceptualization, Data Curation, Formal Analysis, Investigation, Methodology, Software, Validation, Visualization, Writing - Review & Editing. **Zelda Timmel:** Conceptualization, Data Curation, Formal Analysis, Investigation, Software, Validation, Visualization, Writing - Review & Editing. **Christophe Eloy:** Conceptualization, Formal Analysis, Methodology, Supervision, Writing - Review & Editing. **David Robbe:** Conceptualization, Data Curation, Funding Acquisition, Methodology, Project Administration, Resources, Supervision, Writing - Original Draft, Writing - Review & Editing.

## Supplementary figures

**Figure S1:**
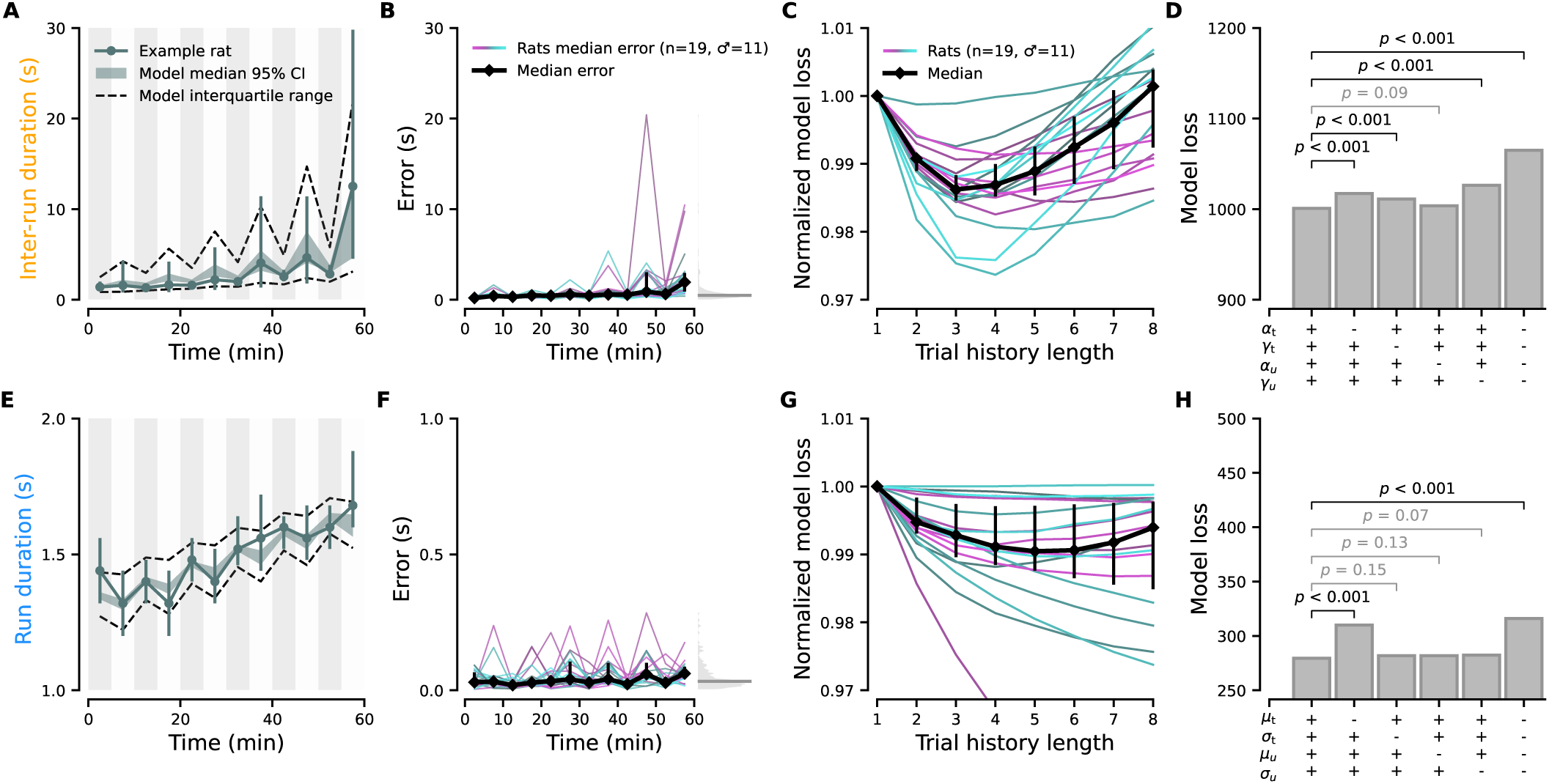
A simple model accurately reconstructs inter-run and run duration dynamics. **A**, Block-by-block median inter-run durations for all the sessions performed in the 94 cm distance condition for one example animal (solid teal line, error bars represent interquartile range). The teal shaded area shows the 95% confidence interval of the synthetic data medians generated using the fitted parameters (*n* = 10000 iterations). Interquartile range (black dashed lines) of all synthetic data generated is also shown. **B**, Block-by-block median error (absolute difference between median experimental data and median synthetic data) for each rat, same condition as A (*n* = 19, thin colored lines). Black, median; error bars, interquartile range. Right inset, distribution of errors pooled across rats and blocks (gray shading, thin horizontal line shows grand median error ∼0.6 seconds). **C**, Total loss of the model when considering the outcome (rewarded or unrewarded) of an increasing number of previous runs (see Methods). For each rat, the loss is normalized by the loss of the model when only the immediately previous outcome is considered. Inter-run durations from all the sessions were analyzed together, irrespective of the action cost condition. **D**, Model loss following parameter ablation. The first bar shows the score of the complete model (α_t_, γ_t_, α_u_ and γ_u_ free, ++++), the last bar shows the score of the fully ablated model (α_t_, γ_t_, α_u_ and γ_u_ fixed to 0,). The other columns have one of the parameters fixed to 0. The significance of the change in the accuracy of the model is determined using a likelihood ratio test. Bonferroni correction is applied. **E-H**, Same as A-D with the run duration model.

**Figure S2:**
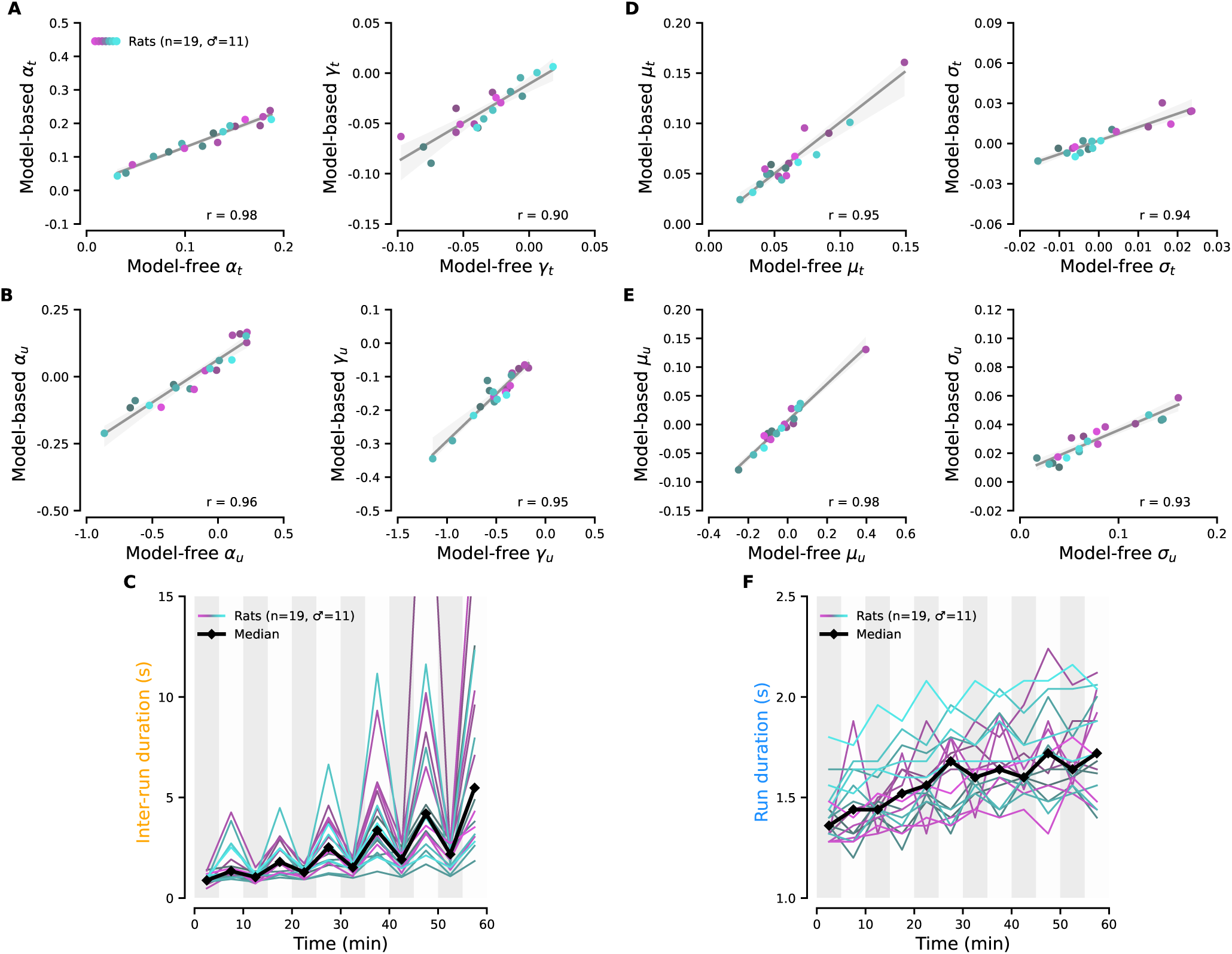
Model-based parameters closely match their model-free counterparts. **A, B**, Correlation across animals between model-derived parameters describing the modulation of inter-run durations by time **(A)** and unrewarded history **(B)**, and their corresponding model-free estimates (Methods). All parameters show strong agreement between the two approaches (Pearson *r* ≥ 0.90), indicating that the model captures the same individual differences as the model-free analysis. **C**, Model-free block-by-block median inter-run durations, computed for each rat using data from all sessions performed under the same condition as in Figure 1B (colored lines). Black line indicates the median across rats. **D-F**, Same analysis as in A-C, but for run durations showing similarly high correlations between model-based and model-free parameter estimates (*r* ≥ 0.93).

**Figure S3:**
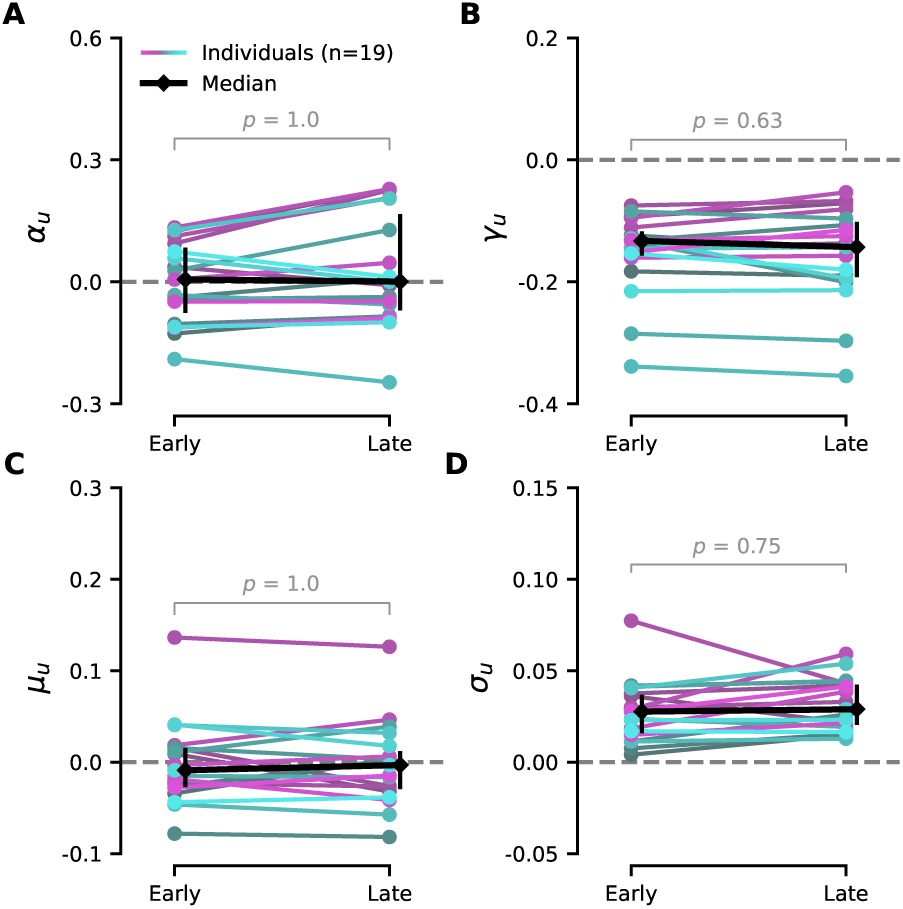
The modulations of model parameters by reward history remained stable throughout the session. **A-D**, Values of α_u_ **(A)**, γ_u_ **(B)**, µ_u_ **(C)**, and σ_u_ **(D)** estimated from the first (early) and last (late) 30 min of the session for each animal (*n* = 19, colored lines). Black dots and error bars show group median and interquartile range. *p*-values from paired permutation tests on median differences are indicated in each panel (Methods).

**Figure S4:**
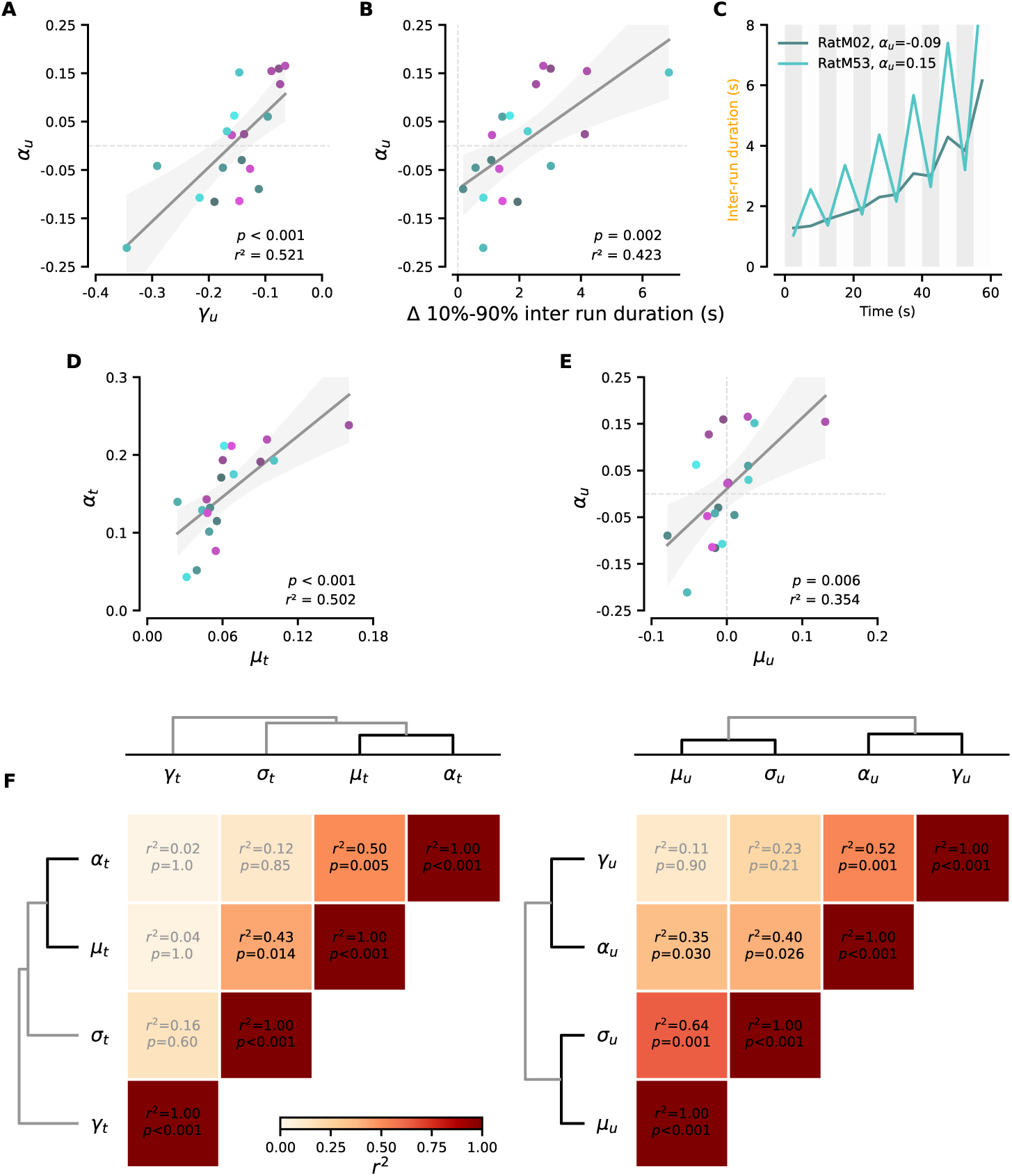
Pairwise correlations between model parameters reveal partial coupling between run timing and speed. **A**, Correlation across animals (*n* = 19) between model-derived α_u_ and γ_u_ for the condition with the treadmill belt immobile and distance set to 94 cm. *p*-values are obtained using a permutation procedure and defined as the fraction of shuffles yielding a correlation coefficient greater in absolute value than the observed one. The shaded area shows the 95% confidence interval of the regression. Pearson correlation coefficients are indicated in each panel (Methods). **B**, Similar to A for α_u_ and the median inter run duration difference between high and low reward probability blocks (Methods). **C**, Block by block inter-run duration median reconstructed from α(*t*, *u*) and γ(*t*, *u*) (Methods) for the two animals at the extremes of the correlation shown in B (the animal with the highest and the animal with the lowest Δ inter-run duration modulation among the 19 animals). The values of α_u_ are indicated in the legend. **D, E**, Similar to A for α_t_ and µ_t_ **(D)** and for α_u_ and µ_u_ **(E)**. **F**, Pairwise correlation matrices and hierarchical dendrograms between the fitted model parameters capturing the modulation of run and inter run duration with time (left) and unrewarded history (right). Pearson correlation coefficients and *p*-values are reported in each cell, with Bonferroni correction applied across all comparisons.

**Figure S5:**
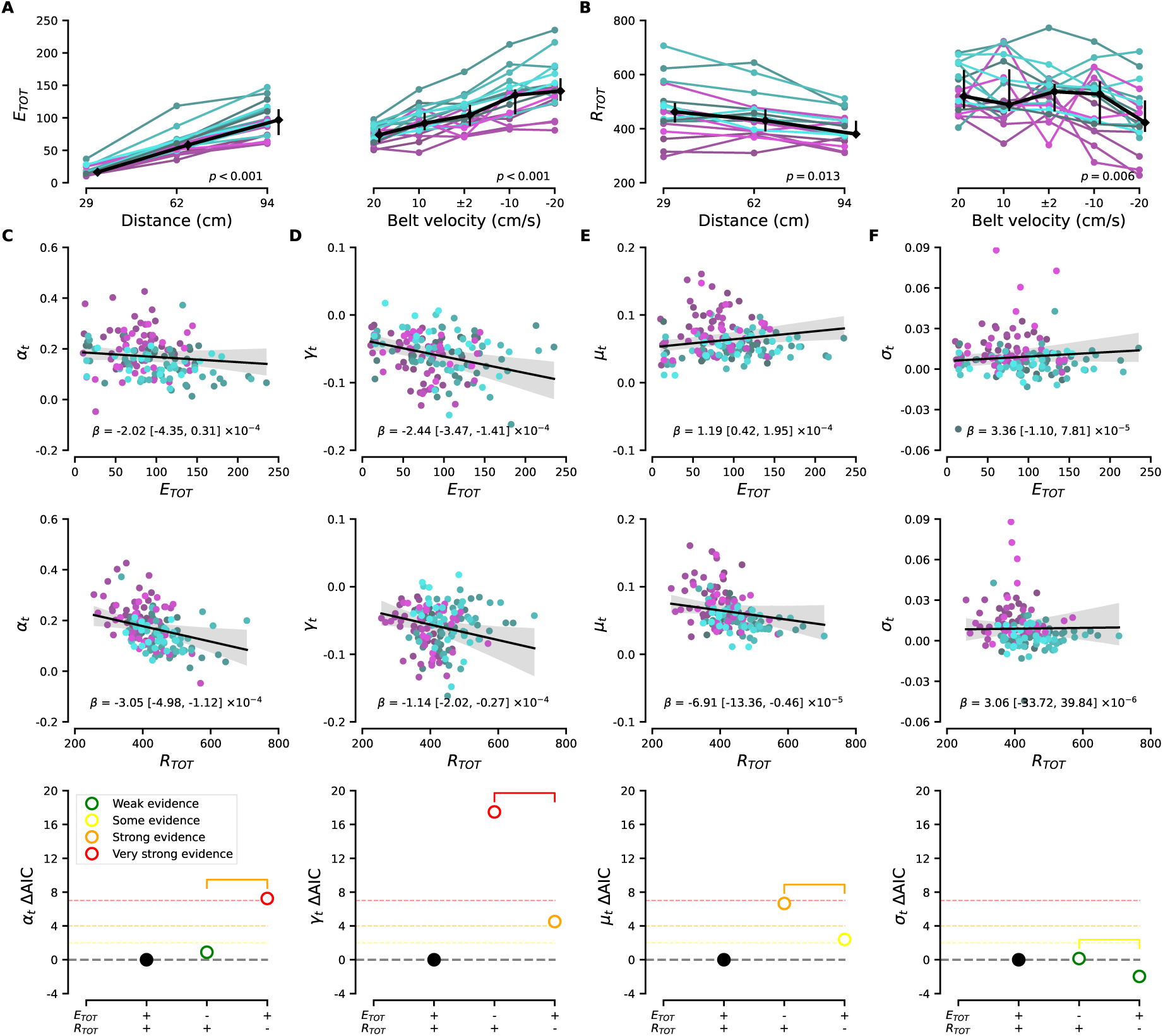
Total effort expenditure and reward collection differentially account for change in inter-run and run duration model parameters over session time. **A**, Estimation of the total amount of effort expended (E_TOT_) by each rat across manipulations of treadmill length and distances (Methods). Each colored line represents the median estimated effort of all sessions for each rat and each cost condition (*n* = 19). Black line and error bars show population median and interquartile range. Slopes were tested using a permutation procedure (Methods); *p*-values indicate the fraction of shuffles with a slope greater than the observed population slope. **B**, Total number of drops collected (R_TOT_) by each rat across manipulation of distance and treadmill belt velocity. Same representation and statistical procedures as A. **C**, Correlation analysis using linear mixed models between α_t_ and the total effort expended (top) or the total number of rewards (middle). Each dot represents one session; the fixed-effect slope β and its 95% confidence interval are indicated. Bottom, ΔAIC comparing the full model against reduced models retaining only (E_TOT_) or only (R_TOT_), computed relative to the full model and interpreted following Burnham and Anderson ([109], see Methods for details). **D-F**, Similar to C for γ_t_ **(D)**, µ_t_ **(E)** and σ_t_ **(F)**.

**Figure S6:**
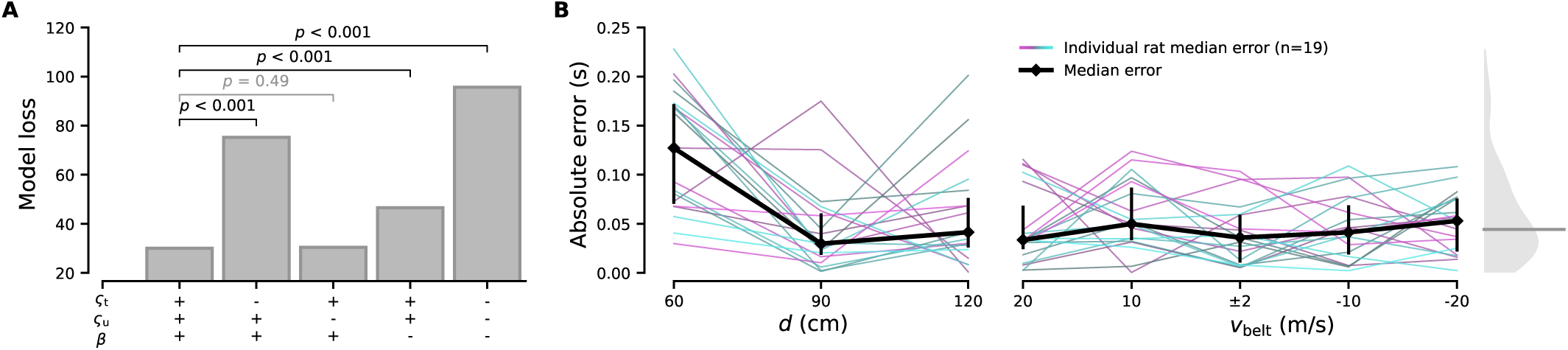
Goodness-of-fit and parameter contribution of the cost-minimization model. **A**, Model loss following parameter ablation. The first bar shows the score of the complete model (ς_t_, ς_u_ and β free, +++), the last bar shows the score of the fully ablated model (—). The significance of the change in the accuracy of the model is determined using a likelihood ratio test with Bonferroni correction (Methods). **B**, Absolute error between median experimental and median synthetic run durations for each rat across distance (*d*, left) and belt velocity (*v*_belt_, right) (*n* = 19, thin colored lines). Black line and error bars show population median and interquartile range. Right inset, distribution of errors pooled across rats and conditions (gray shading, thin horizontal line shows grand median error ∼0.044 seconds).

**Figure S7:**
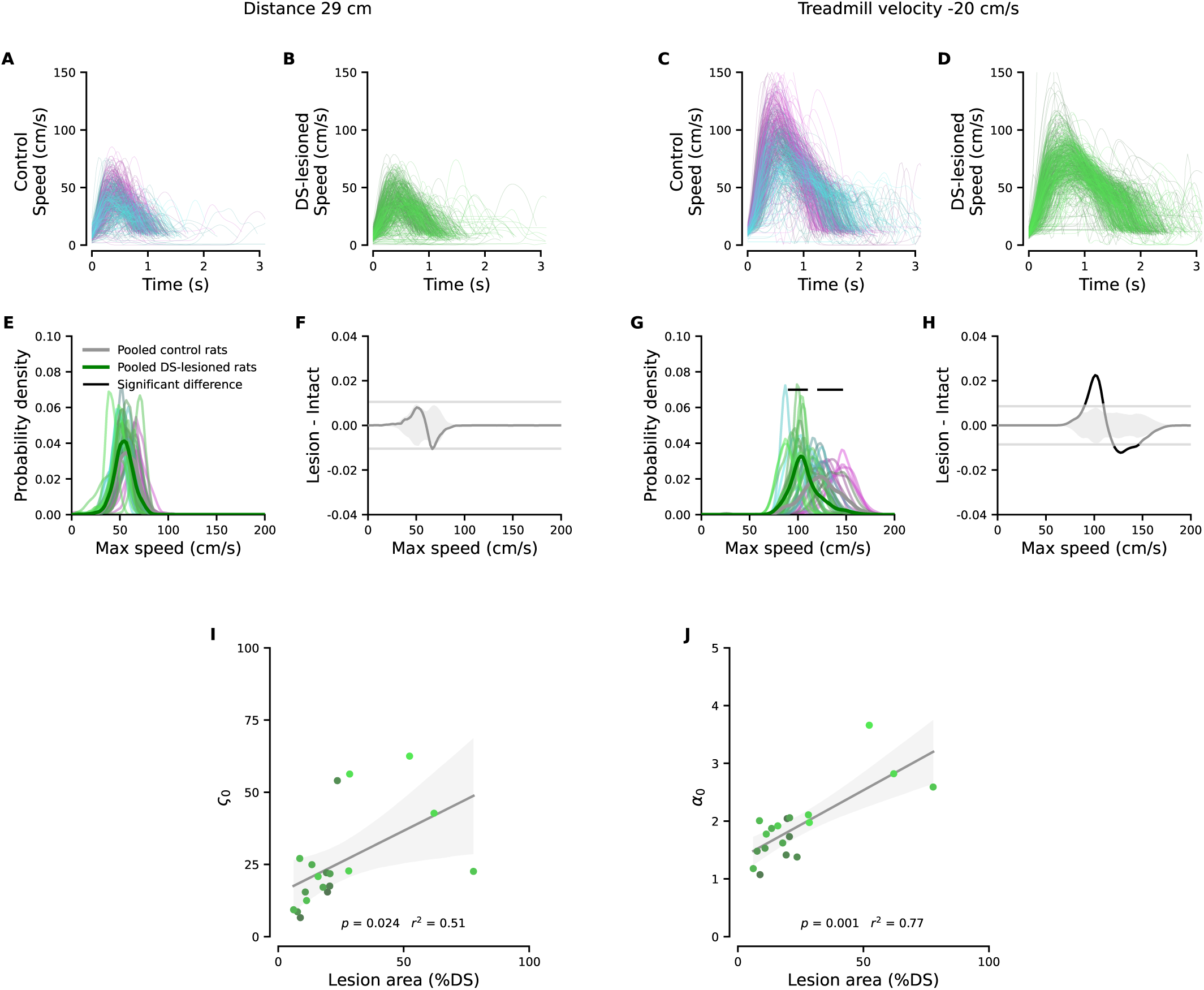
DS-lesion effect on speed profiles and correlation of the lesion extent with model parameters. **A, B**, Individual run speed profiles for control (A; *n* = 19) and DS-lesioned (B; *n* = 19) rats under the lowest motor cost condition (distance 29 cm, treadmill belt stationary), 5 runs per session were randomly shown for each animal. **C, D**, Same as A-B but for the highest motor cost condition (distance 94 cm, treadmill belt velocity = −20 cm/s). **E, F**, Distribution of peak running speeds for control and DS-lesioned animals under the lowest motor cost condition. Left, individual (light traces) and group-average (bold traces) speed distributions. Significant differences between group medians were assessed using a permutation test (black horizontal lines, Methods). Right, difference in probability density between DS-lesioned and control animals (Lesion − Control), with dashed lines indicating the global 95 % confidence interval. **G, H**, Same as in E-F, but for the highest motor cost condition. **I, J**, Correlation between relative sensitivity to movement and time cost (I, ς_0_) or threshold parameter (J, α_0_) and lesion area. Gray shading indicates the 95 % confidence interval of the linear fit. *p*-value is obtained using a shuffling procedure and is defined as the fraction of shuffles with a greater slope than the observed slope.

**Figure S8:**
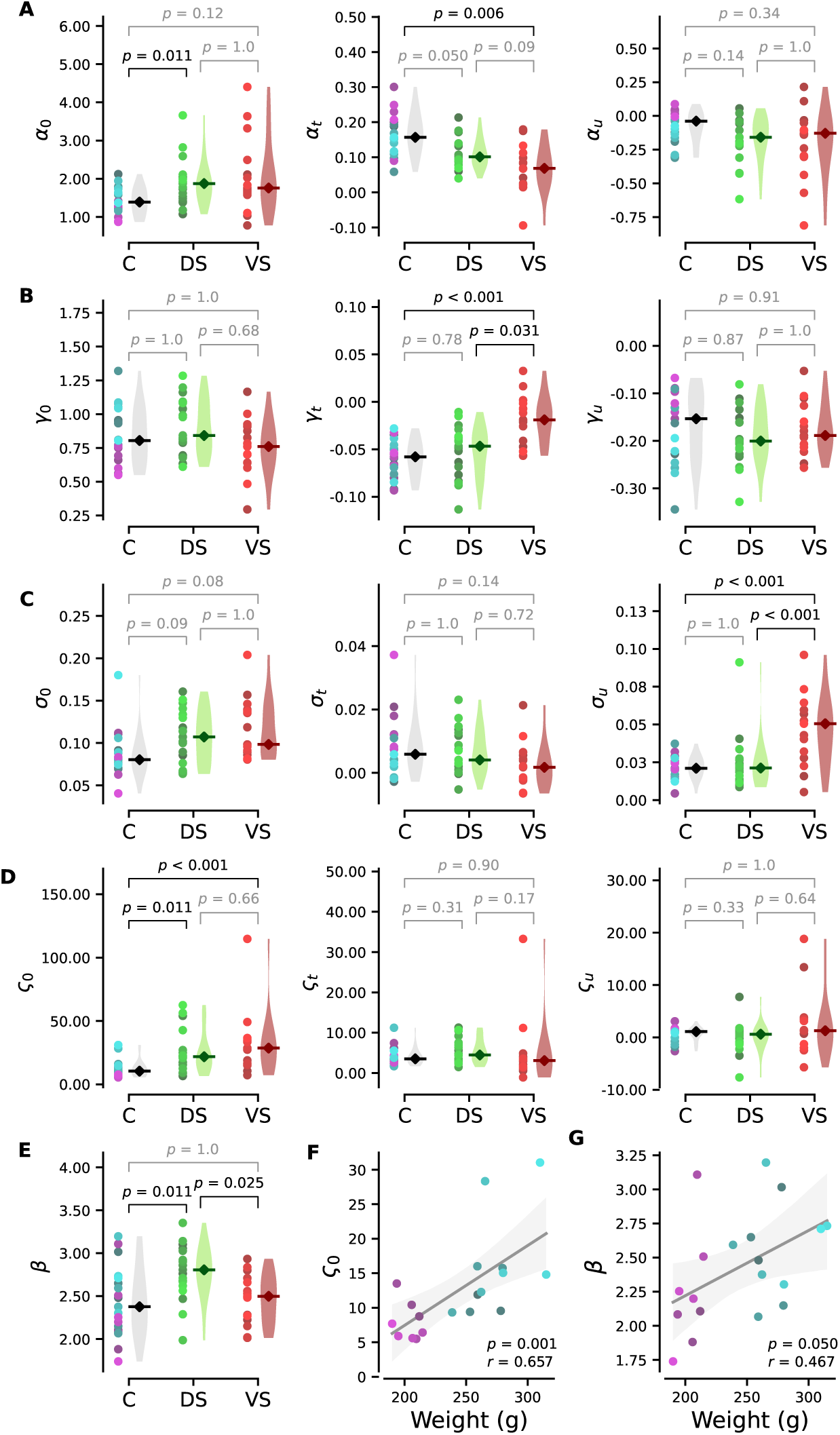
Effect of lesions on model parameters and body-weight dependence of cost minimization parameters. **A**, Inter-group differences in the median α parameter of the inter-run model: baseline value α_0_ (left), modulation with time α_t_ (middle) and modulation with unrewarded history α_u_ (right). Each dot represents one animal; black diamonds and violin plots show group median and distribution for controls (C, *n* = 19), DS-lesioned (DS, *n* = 19), and VS-lesioned (VS, *n* = 15) rats. *p*-values are obtained from a permutation procedure and defined as the fraction of shuffles yielding a difference in group medians greater than or equal to the observed difference. Bonferroni correction is applied (Methods). **B-E**, Similar to A with the γ parameter of the inter-run model, the σ parameter of the run model, the ς and the β parameters of the effort model. **F, G**, Correlation between body weight and the cost minimization model parameters ς (F) and β (G) in control animals (See Methods for details).

### Supplementary video

**Supplementary Video. Preserved crossing duration despite opposing belt velocity.** Two runs performed by the same animal while the treadmill belt speed is set to +20 cm/s (top) or −20 cm/s (bottom). The video is available at: https://drive.google.com/file/d/10NQiavweGwfXNvQAcCj1F4PdQI2pN7bx/view

## References

[1] Niv, Y., Daw, N. D., Joel, D. & Dayan, P. Tonic dopamine: opportunity costs and the control of response vigor. Psychopharmacology 191, 507–520 (2007). Publisher: Springer.

[2] Carland, M. A., Thura, D. & Cisek, P. The Urge to Decide and Act: Implications for Brain Function and Dysfunction. The Neuroscientist 25, 491–511 (2019). URL http://journals.sagepub.com/doi/10.1177/1073858419841553.

[3] Shadmehr, R., Reppert, T. R., Summerside, E. M., Yoon, T. & Ahmed, A. A. Movement vigor as a reflection of subjective economic utility. Trends in neurosciences 42, 323–336 (2019). Publisher: Elsevier.

[4] Myerson, J. & Green, L. Discounting of delayed rewards: Models of individual choice. Journal of the Experimental Analysis of Behavior 64, 263–276 (1995).

[5] Shadmehr, R., Xivry, J. J. O. d., Xu-Wilson, M. & Shih, T.-Y. Temporal Discounting of Reward and the Cost of Time in Motor Control. Journal of Neuroscience 30, 10507–10516 (2010). URL https://www.jneurosci.org/content/30/31/10507. Publisher: Society for Neuroscience Section: Articles.

[6] Haith, A. M., Reppert, T. R. & Shadmehr, R. Evidence for Hyperbolic Temporal Discounting of Reward in Control of Movements. Journal of Neuroscience 32, 11727–11736 (2012). URL https://www.jneurosci.org/content/32/34/11727. Publisher: Society for Neuroscience Section: Articles.

[7] Rigoux, L. & Guigon, E. A model of reward-and effort-based optimal decision making and motor control. PLoS Comput Biol 8, e1002716 (2012).

[8] Choi, J. E. S., Vaswani, P. A. & Shadmehr, R. Vigor of Movements and the Cost of Time in Decision Making. The Journal of Neuroscience 34, 1212–1223 (2014). URL https://www.jneurosci.org/lookup/doi/10.1523/JNEUROSCI.2798-13.2014.

[9] Berret, B. & Jean, F. Why Don’t We Move Slower? The Value of Time in the Neural Control of Action. Journal of Neuroscience 36, 1056–1070 (2016). URL https://www.jneurosci.org/content/36/4/1056. Publisher: Society for Neuroscience Section: Articles.

[10] Summerside, E. M., Shadmehr, R. & Ahmed, A. A. Vigor of reaching movements: reward discounts the cost of effort. Journal of Neurophysiology 119, 2347–2357 (2018). URL https://www.physiology.org/doi/10.1152/jn.00872.2017.

[11] Reppert, T. R. et al. Movement vigor as a traitlike attribute of individuality. Journal of Neurophysiology 120, 741–757 (2018). URL https://www.physiology.org/doi/10.1152/jn.00033.2018.

[12] Yoon, T., Geary, R. B., Ahmed, A. A. & Shadmehr, R. Control of movement vigor and decision making during foraging. Proceedings of the National Academy of Sciences 115 (2018). URL https://pnas.org/doi/full/10.1073/pnas.1812979115.

[13] Kita, K., Du, Y. & Haith, A. M. Evidence for a common mechanism supporting invigoration of action selection and action execution. Journal of Neurophysiology 130, 238–246 (2023). URL https://journals.physiology.org/doi/abs/10.1152/jn.00510.2022. Publisher: American Physiological Society.

[14] Shadmehr, R. & Ahmed, A. A. Vigor: neuroeconomics of movement control (MIT Press, 2020).

[15] Albin, R. L., Young, A. B. & Penney, J. B. The functional anatomy of basal ganglia disorders. Trends in neurosciences 12, 366–375 (1989).

[16] Hwang, E. J. The basal ganglia, the ideal machinery for the cost-benefit analysis of action plans. Frontiers in neural circuits 7, 121 (2013).

[17] Frank, M. J. Adaptive cost-benefit control fueled by striatal dopamine. Annual Review of Neuroscience 48 (2025).

[18] Mazzoni, P., Hristova, A. & Krakauer, J. W. Why Don’t We Move Faster? Parkinson’s Disease, Movement Vigor, and Implicit Motivation. The Journal of Neuroscience 27, 7105–7116 (2007). URL https://www.jneurosci.org/lookup/doi/10.1523/JNEUROSCI.0264-07.2007.

[19] Baraduc, P., Thobois, S., Gan, J., Broussolle, E. & Desmurget, M. A Common Optimization Principle for Motor Execution in Healthy Subjects and Parkinsonian Patients. The Journal of Neuroscience 33, 665–677 (2013). URL https://www.jneurosci.org/lookup/doi/10.1523/JNEUROSCI.1482-12.2013.

[20] Suzuki, S., Lawlor, V. M., Cooper, J. A., Arulpragasam, A. R. & Treadway, M. T. Distinct regions of the striatum underlying effort, movement initiation and effort discounting. Nature human behaviour 5, 378–388 (2021).

[21] Herz, D. M. & Brown, P. Moving, fast and slow: behavioural insights into bradykinesia in parkinson’s disease. Brain 146, 3576–3586 (2023).

[22] Kable, J. W. & Glimcher, P. W. The neural correlates of subjective value during intertemporal choice. Nature neuroscience 10, 1625–1633 (2007).

[23] Yaakub, S. N. et al. Non-invasive ultrasonic neuromodulation of the human nucleus accumbens impacts reward sensitivity. Nature Communications 16, 10192 (2025).

[24] Pessiglione, M. et al. How the brain translates money into force: a neuroimaging study of subliminal motivation. science 316, 904–906 (2007).

[25] Aberman, J. Nucleus accumbens dopamine depletions make rats more sensitive to high ratio requirements but do not impair primary food reinforcement. Neuroscience 92, 545–552 (1999).

[26] Ishiwari, K., Weber, S. M., Mingote, S., Correa, M. & Salamone, J. D. Accumbens dopamine and the regulation of effort in food-seeking behavior: modulation of work output by different ratio or force requirements. Behavioural Brain Research 151, 83–91 (2004). URL https://www.sciencedirect.com/science/article/pii/S0166432803002924.

[27] Wang, A. Y., Miura, K. & Uchida, N. The dorsomedial striatum encodes net expected return, critical for energizing performance vigor. Nature Neuroscience 16, 639–647 (2013). URL https://www.nature.com/articles/nn.3377.

[28] R ueda-Orozco, P. E. & Robbe, D. The striatum multiplexes contextual and kinematic information to constrain motor habits execution. Nature Neuroscience 18, 453–460 (2015). URL 10.1038/nn.3924.

[29] Panigrahi, B. et al. Dopamine Is Required for the Neural Representation and Control of Movement Vigor. Cell 162, 1418–1430 (2015). URL https://linkinghub.elsevier.com/retrieve/pii/S0092867415010272.

[30] Bartholomew, R. A. et al. Striatonigral control of movement velocity in mice. European Journal of Neuroscience 43, 1097–1110 (2016).

[31] Hamid, A. A. et al. Mesolimbic dopamine signals the value of work. Nature Neuroscience 19, 117–126 (2016). URL 10.1038/nn.4173.

[32] Yttri, E. A. & Dudman, J. T. Opponent and bidirectional control of movement velocity in the basal ganglia. Nature 533, 402–406 (2016). URL 10.1038/nature17639.

[33] Sales-Carbonell, C. et al. No discrete start/stop signals in the dorsal striatum of mice performing a learned action. Current Biology 28, 3044–3055.e5 (2018). URL 10.1016/j.cub.2018.07.038.

[34] Mohebi, A. et al. Dissociable dopamine dynamics for learning and motivation. Nature 570, 65–70 (2019). URL 10.1038/s41586-019-1235-y.

[35] Jurado-Parras, M.-T. et al. The dorsal striatum energizes motor routines. Current Biology 30, 4362–4372.e6 (2020). URL 10.1016/j.cub.2020.08.049.

[36] Fobbs, W. C. et al. Continuous representations of speed by striatal medium spiny neurons. Journal of Neuroscience 40, 1679–1688 (2020).

[37] Mizes, K. G. C., Lindsey, J., Escola, G. S. & Ölveczky, B. P. Dissociating the contributions of sensorimotor striatum to automatic and visually guided motor sequences. Nature Neuroscience 26, 1791–1804 (2023). URL 10.1038/s41593-023-01431-3.

[38] Levcik, D. et al. Nucleus accumbens shell neurons encode the kinematics of reward approach locomotion. Neuroscience 524, 181–196 (2023).

[39] Beas, S. et al. Dissociable encoding of motivated behavior by parallel thalamo-striatal projections. Current Biology 34, 1549–1560 (2024).

[40] Zheng, Q. et al. The role of striatum in controlling waiting during reactive and self-timed behaviors. Journal of Neuroscience 45 (2025).

[41] Shoemaker, C. T. et al. A2a-positive neurons in the nucleus accumbens core regulate effort exertion. Journal of Neuroscience 45 (2025).

[42] Packard, M. G. & Knowlton, B. J. Learning and memory functions of the basal ganglia. Annual Review of Neuroscience 25, 563–593 (2002).

[43] Yin, H. H. & Knowlton, B. J. The role of the basal ganglia in habit formation. Nature Reviews Neuroscience 7, 464–476 (2006).

[44] Balleine, B. W. & O’Doherty, J. P. Human and rodent homologs in action control: corticostriatal determinants of goal-directed and habitual action. Neuropsychopharmacology 35, 48–69 (2010).

[45] Graybiel, A. M. & Grafton, S. T. The striatum: where skills and habits meet. Cold Spring Harbor perspectives in biology 7, a021691 (2015).

[46] Arber, S. & Costa, R. M. Networking brainstem and basal ganglia circuits for movement. Nature Reviews Neuroscience 23, 342–360 (2022).

[47] Mogenson, G. J., Jones, D. L. & Yim, C. Y. From motivation to action: functional interface between the limbic system and the motor system. Progress in Neurobiology 14, 69–97 (1980).

[48] Cardinal, R. N., Parkinson, J. A., Hall, J. & Everitt, B. J. Emotion and motivation: the role of the amygdala, ventral striatum, and prefrontal cortex. Neuroscience & Biobehavioral Reviews 26, 321–352 (2002).

[49] Kelley, A. E. Ventral striatal control of appetitive motivation: role in ingestive behavior and reward-related learning. Neuroscience & biobehavioral reviews 27, 765–776 (2004).

[50] Berridge, K. C. The debate over dopamine’s role in reward: the case for incentive salience. Psychopharmacology 191, 391–431 (2007).

[51] Haber, S. N. & Knutson, B. The reward circuit: linking primate anatomy and human imaging. Neuropsychopharmacology 35, 4–26 (2010).

[52] Turner, R. S. & Desmurget, M. Basal ganglia contributions to motor control: a vigorous tutor. Current Opinion in Neurobiology 20, 704–716 (2010). URL https://linkinghub.elsevier.com/retrieve/pii/S095943881000142X.

[53] Dudman, J. T. & Krakauer, J. W. The basal ganglia: from motor commands to the control of vigor. Current Opinion in Neurobiology 37, 158–166 (2016). URL https://www.sciencedirect.com/science/article/pii/S095943881630006X. Neurobiology of cognitive behavior.

[54] Thura, D., Haith, A. M., Derosiere, G. & Duque, J. The integrated control of decision and movement vigor. Trends in Cognitive Sciences (2025).

[55] Heathcote, A. Fitting wald and ex-wald distributions to response time data: An example using functions for the s-plus package. Behavior Research Methods, Instruments, & Computers 36, 678–694 (2004).

[56] Ratcliff, R. & Strayer, D. Modeling simple driving tasks with a one-boundary diffusion model. Psychonomic Bulletin & Review 21, 577–589 (2014). URL 10.3758/s13423-013-0541-x.

[57] Ratcliff, R. Modeling one-choice and two-choice driving tasks. *Attention, Perception*, & Psychophysics 77, 2134–2144 (2015). URL 10.3758/s13414-015-0911-8.

[58] Steingroever, H., Wabersich, D. & Wagenmakers, E.-J. Modeling across-trial variability in the wald drift rate parameter. Behavior Research Methods 53, 1060–1076 (2021).

[59] Pekny, S. E., Izawa, J. & Shadmehr, R. Reward-dependent modulation of movement variability. Journal of Neuroscience 35, 4015–4024 (2015).

60. Schaffhauser, M., et al. A novel foraging task reveals cognitive and motor processes underlying behavioral flexibility. bioRxiv (2025). URL https://www.biorxiv.org/content/early/2025/10/28/2025.09.09.675194. https://www.biorxiv.org/content/early/2025/10/28/2025.09.09.675194.full.pdf.

[61] Amsel, A. The role of frustrative nonreward in noncontinuous reward situations. Psychological bulletin 55, 102 (1958).

[62] Papini, M. R., Guarino, S., Hagen, C. & Torres, C. Incentive disengagement and the adaptive significance of frustrative nonreward. Learning & behavior 50, 372–388 (2022).

[63] Todorov, E. Optimality principles in sensorimotor control. Nature neuroscience 7, 907–915 (2004).

64. Verdel, D., Bruneau, O., Sahm, G., Vignais, N. & Berret, B. The value of time in the invigoration of human movements when interacting with a robotic exoskeleton. Science Advances 9, eadh9533 (2023). URL https://www.science.org/doi/abs/10.1126/sciadv.adh9533. https://www.science.org/doi/pdf/10.1126/sciadv.adh9533.

[65] Ralston, H. J. Energy-speed relation and optimal speed during level walking. Internationale Zeitschrift für angewandte Physiologie einschließlich Arbeitsphysiologie 17, 277–283 (1958). URL 10.1007/BF00698754.

[66] Alexander, R. M. Principles of animal locomotion (Princeton university press, 2003).

[67] Taylor, C. R., Heglund, N. C. & Maloiy, G. M. Energetics and mechanics of terrestrial locomotion. i. metabolic energy consumption as a function of speed and body size in birds and mammals. Journal of Experimental Biology 97, 1–21 (1982).

[68] Härmson, O. et al. Hierarchical encoding of reward, effort and choice across the cortex and basal ganglia during cost-benefit decision making. bioRxiv 2023–10 (2023).

[69] Eshel, N. et al. Striatal dopamine integrates cost, benefit, and motivation. Neuron S0896–6273(23)00843–7 (2023).

[70] Bartra, O., McGuire, J. T. & Kable, J. W. The valuation system: a coordinate-based meta-analysis of bold fmri experiments examining neural correlates of subjective value. Neuroimage 76, 412–427 (2013).

[71] Thura, D., Cos, I., Trung, J. & Cisek, P. Context-Dependent Urgency Influences Speed–Accuracy Trade-Offs in Decision-Making and Movement Execution. Journal of Neuroscience 34, 16442–16454 (2014). URL https://www.jneurosci.org/content/34/49/16442. Publisher: Society for Neuroscience Section: Articles.

[72] Reynaud, A. J., Saleri Lunazzi, C. & Thura, D. Humans sacrifice decision-making for action execution when a demanding control of movement is required. Journal of Neurophysiology 124, 497–509 (2020). URL https://journals.physiology.org/doi/full/10.1152/jn.00220.2020. Publisher: American Physiological Society.

73. Fievez, F., et al. Task goals shape the relationship between decision and movement speed. bioRxiv (2023). URL https://www.biorxiv.org/content/early/2023/12/30/2023.12.29.573524. https://www.biorxiv.org/content/early/2023/12/30/2023.12.29.573524.full.pdf.

[74] Saleri, C. & Thura, D. Evidence for interacting but decoupled controls of decisions and movements in nonhuman primates. Journal of Neurophysiology 132, 1470–1480 (2024).

[75] Conessa, A., Boutin, A. & Berret, B. Evidence for separate processes underlying movement and decision vigor in a reward-oriented task. bioRxiv 2025–11 (2025).

[76] Berret, B. & Baud-Bovy, G. Evidence for a cost of time in the invigoration of isometric reaching movements. Journal of Neurophysiology 127, 689–701 (2022). URL https://journals.physiology.org/doi/10.1152/jn.00536.2021.

[77] Shadmehr, R., Huang, H. J. & Ahmed, A. A. A representation of effort in decision-making and motor control. Current biology 26, 1929–1934 (2016).

[78] Polat, L., Harpaz, T. & Zaidel, A. Rats rely on airflow cues for self-motion perception. Current Biology 34, 4248–4260 (2024).

[79] Madhav, M. S. et al. Control and recalibration of path integration in place cells using optic flow. Nature neuroscience 27, 1599–1608 (2024).

[80] Ludwig, C. J. et al. The influence of visual flow and perceptual load on locomotion speed. Attention, Perception, & Psychophysics 80, 69–81 (2018).

[81] Salamone, J. D. & Correa, M. The mysterious motivational functions of mesolimbic dopamine. Neuron 76, 470–485 (2012). Publisher: Elsevier.

[82] Yin, H. H. The basal ganglia in action. The Neuroscientist 23, 299–313 (2017).

[83] Berke, J. D. What does dopamine mean? Nature Neuroscience 21, 787–793 (2018). URL 10.1038/s41593-018-0152-y.

[84] Yttri, E. A. & Dudman, J. T. A proposed circuit computation in basal ganglia: History-dependent gain. Movement Disorders 33, 704–716 (2018).

[85] Robbe, D. & Dudman, J. T. The Basal Ganglia Invigorate Actions and Decisions. In The Cognitive Neurosciences (The MIT Press, 2020). URL 10.7551/mitpress/11442.003.0058. https://direct.mit.edu/book/chapter-pdf/2053718/c037600_9780262356176.pdf.

[86] Tan, B. et al. Dynamic processing of hunger and thirst by common mesolimbic neural ensembles. Proceedings of the National Academy of Sciences of the United States of America 119, e2211688119 (2022). URL https://pmc.ncbi.nlm.nih.gov/articles/PMC9618039/.36252036.

[87] Aitken, T. J., Greenfield, V. Y. & Wassum, K. M. Nucleus accumbens core dopamine signaling tracks the need-based motivational value of food-paired cues. Journal of neurochemistry 136, 1026–1036 (2016). URL https://pmc.ncbi.nlm.nih.gov/articles/PMC4819964/.26715366.

[88] Castro, D. C., Cole, S. L. & Berridge, K. C. Lateral hypothalamus, nucleus accumbens, and ventral pallidum roles in eating and hunger: Interactions between homeostatic and reward circuitry. Frontiers in Systems Neuroscience 9, 90 (2015). 26124708.

[89] Müller, T., Klein-Flügge, M. C., Manohar, S. G., Husain, M. & Apps, M. A. Neural and computational mechanisms of momentary fatigue and persistence in effort-based choice. Nature Communications 12, 4593 (2021).

[90] Banwinkler, M. et al. Putaminal dopamine modulates movement motivation in parkinson’s disease. bioRxiv 2024–03 (2024).

[91] Kravitz, A. V. et al. Regulation of parkinsonian motor behaviours by optogenetic control of basal ganglia circuitry. Nature 466, 622–626 (2010).

[92] Roseberry, T. K. et al. Cell-type-specific control of brainstem locomotor circuits by basal ganglia. Cell 164, 526–537 (2016).

[93] Cregg, J. M., Sidhu, S. K., Leiras, R. & Kiehn, O. Basal ganglia–spinal cord pathway that commands locomotor gait asymmetries in mice. Nature Neuroscience 27, 716–727 (2024). URL https://www.nature.com/articles/s41593-024-01569-8.

[94] Hardcastle, K. et al. Differential kinematic coding in sensorimotor striatum across species-typical and learned behaviors reflects a difference in control. bioRxiv 2023–10 (2023).

[95] Kiehn, O. Decoding the organization of spinal circuits that control locomotion. Nature Reviews Neuroscience 17, 224–238 (2016).

[96] Walton, M., Kennerley, S. W., Bannerman, D., Phillips, P. & Rushworth, M. F. Weighing up the benefits of work: behavioral and neural analyses of effort-related decision making. Neural networks 19, 1302–1314 (2006).

[97] Hintiryan, H. et al. The mouse cortico-striatal projectome. Nature neuroscience 19, 1100–1114 (2016).

[98] Kurniawan, I. T. et al. Choosing to make an effort: the role of striatum in signaling physical effort of a chosen action. Journal of neurophysiology 104, 313–321 (2010).

[99] Mondragón-González, S. L., Schreiweis, C. & Burguière, E. Closed-loop recruitment of striatal interneurons prevents compulsive-like grooming behaviors. Nature neuroscience 27, 1148–1156 (2024).

[100] Geramita, M. A., Ahmari, S. E. & Yttri, E. A. The striatal indirect pathway mediates hesitation. Nature Neuroscience 29, 287–292 (2026).

[101] Reinagel, P. Training rats using water rewards without water restriction. Frontiers in Behavioral Neuroscience 12 (2018). URL https://www.frontiersin.org/articles/10.3389/fnbeh.2018.00084.

[102] Fränti, P. & Mariescu-Istodor, R. Averaging gps segments competition 2019. Pattern Recognition 112, 107730 (2021). URL https://www.sciencedirect.com/science/article/pii/S0031320320305331.

[103] Anders, R., Alario, F. & Van Maanen, L. The shifted Wald distribution for response time data analysis. Psychological methods 21, 309 (2016). URL https://psycnet.apa.org/journals/met/21/3/309/. Publisher: American Psychological Association.

[104] Ratcliff, R. & Van Dongen, H. P. A. Diffusion model for one-choice reaction-time tasks and the cognitive effects of sleep deprivation. Proceedings of the National Academy of Sciences 108, 11285–11290 (2011). URL https://pnas.org/doi/full/10.1073/pnas.1100483108.

[105] Virtanen, P. et al. SciPy 1.0: Fundamental Algorithms for Scientific Computing in Python. Nature Methods 17, 261–272 (2020).

[106] Seabold, S. & Perktold, J. statsmodels: Econometric and statistical modeling with python. In 9th Python in Science Conference (2010).

[107] Nakagawa, S. & Schielzeth, H. A general and simple method for obtaining r2 from generalized linear mixed-effects models. Methods in Ecology and Evolution 4, 133–142 (2013). URL https://besjournals.onlinelibrary.wiley.com/doi/abs/10.1111/j.2041-210x.2012.00261.x. https://besjournals.onlinelibrary.wiley.com/doi/pdf/10.1111/j.2041-210x.2012.00261.x.

[108] Pedregosa, F. et al. Scikit-learn: Machine learning in Python. Journal of Machine Learning Research 12, 2825–2830 (2011).

[109] Burnham, K. P. & Anderson, D. R. Multimodel inference: understanding aic and bic in model selection. Sociological methods & research 33, 261–304 (2004).

[110] Schmidt-Nielsen, K. Locomotion: energy cost of swimming, flying, and running. Science 177, 222–228 (1972).

[111] Alexander, R. M. Models and the scaling of energy costs for locomotion. Journal of Experimental Biology 208, 1645–1652 (2005).

[112] Busse, S. & Schwarting, R. K. Decoupling actions from consequences: Dorsal hippocampal lesions facilitate instrumental performance, but impair behavioral flexibility in rats. Frontiers in behavioral neuroscience 10, 118 (2016).

[113] Schindelin, J., et al. Fiji: an open-source platform for biological-image analysis. Nat Meth 9, 676–682 (2012). URL 10.1038/nmeth.2019.

